# Large populations from few founders: founder-aware likelihoods for inferring cell proliferation rates

**DOI:** 10.64898/2026.11.20.767990

**Authors:** Lifeng Han, Marley M. Assefa

## Abstract

Gaussian approximations to demographic noise are usually justified by large numbers of cells. In populations described by branching processes, however, a large population descended from few founders is not approximately Gaussian. A culture grown from few cells can contain thousands of cells yet retain large, skewed variation from its earliest divisions, deaths, and phenotypic switches. A random establishment amplitude, the limit of the population size divided by its exponential growth factor, summarizes this variation. We propose a founder-aware likelihood for cell cultures observed at an endpoint. Given the establishment amplitude, it approximates the remaining demographic fluctuations by a Gaussian distribution and then averages over the distribution of the amplitude. Its conditional Gaussian approximation requires a large population rather than many founders, and the likelihood approaches a single Gaussian distribution as the founder count grows. We compare it with a population Gaussian likelihood developed for many-founder cultures, using a pure-birth (Yule) process, whose cell-count distribution is known exactly, and exact simulations of a two-type process with division, death, and reversible switching. With one founder per culture, the founder-aware likelihood estimates rates more accurately than the population Gaussian likelihood, and its confidence sets are longer but have coverage close to the nominal level. With 50 founders per culture, the two likelihoods perform comparably. Founder-aware likelihoods thus suit clonal and other few-founder experiments.

## 1 Introduction

Rates of cell division are commonly estimated from replicate cultures whose cells are counted, often by phenotypic state, after a period of growth (Gupta et al. 2011; Gunnarsson et al. 2023; Wu et al. 2024). In heterogeneous populations, cells also switch between phenotypic states, so the growth of each state reflects its own division and death as well as the exchange of cells between states (Sharma et al. 2010; Gupta et al. 2011; Shaffer et al. 2017; Russo et al. 2022). Branching models represent division, death, and switching at the level of individual cells and connect these mechanisms to both mean counts and demographic variability (Kimmel and Axelrod 2015; Durrett 2015; Hyrien et al. 2010; Gunnarsson et al. 2020, 2023). When rates and their uncertainty are inferred from replicate cultures, the statistical approximation used for this demographic variability affects both the estimates and their reported uncertainty. The choice of approximation matters most when cultures start from very few cells, as in colony-formation assays (Franken et al. 2006), clonal assays, and lineage-barcoding studies that follow hundreds to more than a thousand clones, each founded by one cell (Hyrien et al. 2010; Mathis et al. 2017).

Founder count and observed population size play different roles. A culture seeded with many independent cells contains contributions from many founding lineages. As the founder count increases, a central limit theorem justifies a Gaussian approximation to its composition (Yakovlev and Yanev 2009; Gunnarsson et al. 2023; Wu et al. 2024, 2025). A culture descended from one cell may contain thousands of cells at observation, but its early divisions and switches affect the subsequent population as a whole. Under supercritical branching growth, the population size divided by its exponential growth factor converges to a random limit, the establishment amplitude, which records the effect of these early events on all later population sizes (Athreya and Ney 1972). Growth does not create additional independent founding histories. Consequently, a large endpoint count and a large founder count justify different Gaussian approximations. A large endpoint count makes the fluctuations that remain after the earliest events approximately Gaussian, conditional on the establishment amplitude; only a large founder count makes the amplitude, and hence the endpoint population, approximately Gaussian. For birth–death processes, Davison et al. (2021) found that Gaussian likelihood estimation loses accuracy when populations start from a single individual but performs well when they start from many. For multitype branching processes, Luo et al. (2026, Section 5.1) found that the Gaussian approximation of Gunnarsson et al. (2023) represented the distribution of population sizes much less accurately than their saddlepoint approximation, and attributed the difference to its reliance on large initial populations.

Independent endpoint designs, in which each culture is measured once, are particularly sensitive to the difference between founder count and population size. A single observation cannot distinguish a favorable early history from faster growth, because both produce a larger culture. Longitudinal observations of one population can partly distinguish these two explanations through the growth between observations. With independent endpoint observations, these explanations can be distinguished only through the distribution of amplitudes across cultures. With few founders, the amplitude distribution therefore becomes part of the information about growth rates.

Inference for single-founder cultures has an established statistical basis. Gaussian pseudo-likelihoods use a Gaussian criterion built from population means and covariances without assuming Gaussian data, and have been applied to single-founder clones observed once (Hyrien et al. 2010). Chen and Hyrien (2011) showed that, with correct moments, identifiability, and suitable regularity conditions, Gaussian pseudo-likelihood estimators are consistent and asymptotically normal as the number of independent cultures increases. They also proposed an exponential pseudo-likelihood based on the limiting distribution of a single-founder pure-birth population. This approximate distribution uses the amplitude distribution but omits the fluctuations around it (Section 2.5). Three different limits are involved in this setting, and each justifies a different approximation:

i. an increasing number of independent cultures, which gives consistency of Gaussian pseudo-likelihood estimators and, for a correctly specified likelihood, chi-square calibration of profile-likelihood confidence sets;
ii. an increasing founder count per culture, which makes a Gaussian distribution accurate for each culture;
iii. an increasing observation time at fixed founder count, which defines the establishment amplitude and motivates the conditional residual approximation of Section 2.3.

Consistency does not imply efficiency. For single-founder birth–death populations observed longitudinally, Chen and Hyrien (2011, Section 4.2 and Table 2) found Gaussian pseudo-likelihood estimates substantially less accurate than exact maximum likelihood estimates, and exact maximum likelihood numerically unstable when populations grew large. For large populations founded by few cells, neither approach is satisfactory. Gaussian pseudo-likelihood is easy to compute but does not represent the non-Gaussian distribution inherited from the founders, whereas the exact likelihood is difficult to evaluate (Ho et al. 2018; Luo et al. 2026).

We propose the founder-aware likelihood, which distinguishes the establishment amplitude from the remaining demographic fluctuations. Conditional on the establishment amplitude, it approximates these fluctuations by a Gaussian distribution. Integrating over the amplitude distribution then gives a mixture likelihood. The mixture retains the non-Gaussian amplitude distribution inherited from the founders, which is not represented by Gaussian pseudo-likelihood. Evaluating the mixture requires a numerical integral over a scalar amplitude for each culture, in place of an exact likelihood calculation. A separate component represents cultures that are empty at observation. The construction applies to multitype Markov branching processes with a known initial state and requires only the mean-growth matrix, the infinitesimal covariance of the branching process, and the numbers of founders of each type.

Our principal comparison is the population Gaussian likelihood of Gunnarsson et al. (2023). This likelihood models each culture’s observation by a Gaussian distribution with the exact population mean and covariance plus measurement error. Gunnarsson et al. (2023) developed it for the model of proliferation and reversible phenotypic switching and the independent endpoint design that we also use. They justified the Gaussian form by a central limit theorem as the number of founders increases. We ask whether retaining the distribution of the establishment amplitude improves the approximation to the data distribution, and the resulting rate estimates and uncertainty sets, relative to the population Gaussian likelihood.

Random amplitude and random time shift are established descriptions of early stochastic growth (Barbour et al. 2015, 2016; Baker et al. 2018; Chigansky et al. 2019; Cheek and Antal 2018). During exponential growth, multiplying a trajectory by a positive amplitude is equivalent to translating time. Morris et al. (2024) developed methods to compute random time-shift distributions, and Morris et al. (2025) integrated a random time shift into a hierarchical likelihood for viral dynamics. Building on this prior work, our contribution is the Gaussian approximation to demographic fluctuations conditional on the amplitude, and a numerical assessment of the resulting mixture likelihood for rate inference.

We test the founder-aware likelihood on the two-type model of proliferation and reversible phenotypic switching studied by Gunnarsson et al. (2023). We compare five likelihoods (Table 1). The first two are the population Gaussian likelihood of Gunnarsson et al. (2023) and an adaptation that represents empty cultures in the same way as the founder-aware likelihood. The remaining three are the founder-aware likelihood and two variants that change only the treatment of demographic variation conditional on amplitude. One variant omits this variation, and the other describes it by a single fitted variance. We first use the Yule process, in which a single cell type divides at a constant rate and the relevant distributions are known exactly, to explain the approximation and identify its limitation at early times. A simulation study of the two-type model then compares rate estimates, profile-likelihood confidence sets, predictive accuracy, and computational cost. With one founder per culture, the founder-aware likelihood estimates rates more accurately than the population Gaussian likelihood, and its confidence sets are longer but have coverage close to the nominal level. With 50 founders, the two likelihoods perform comparably.

**Table 1.**
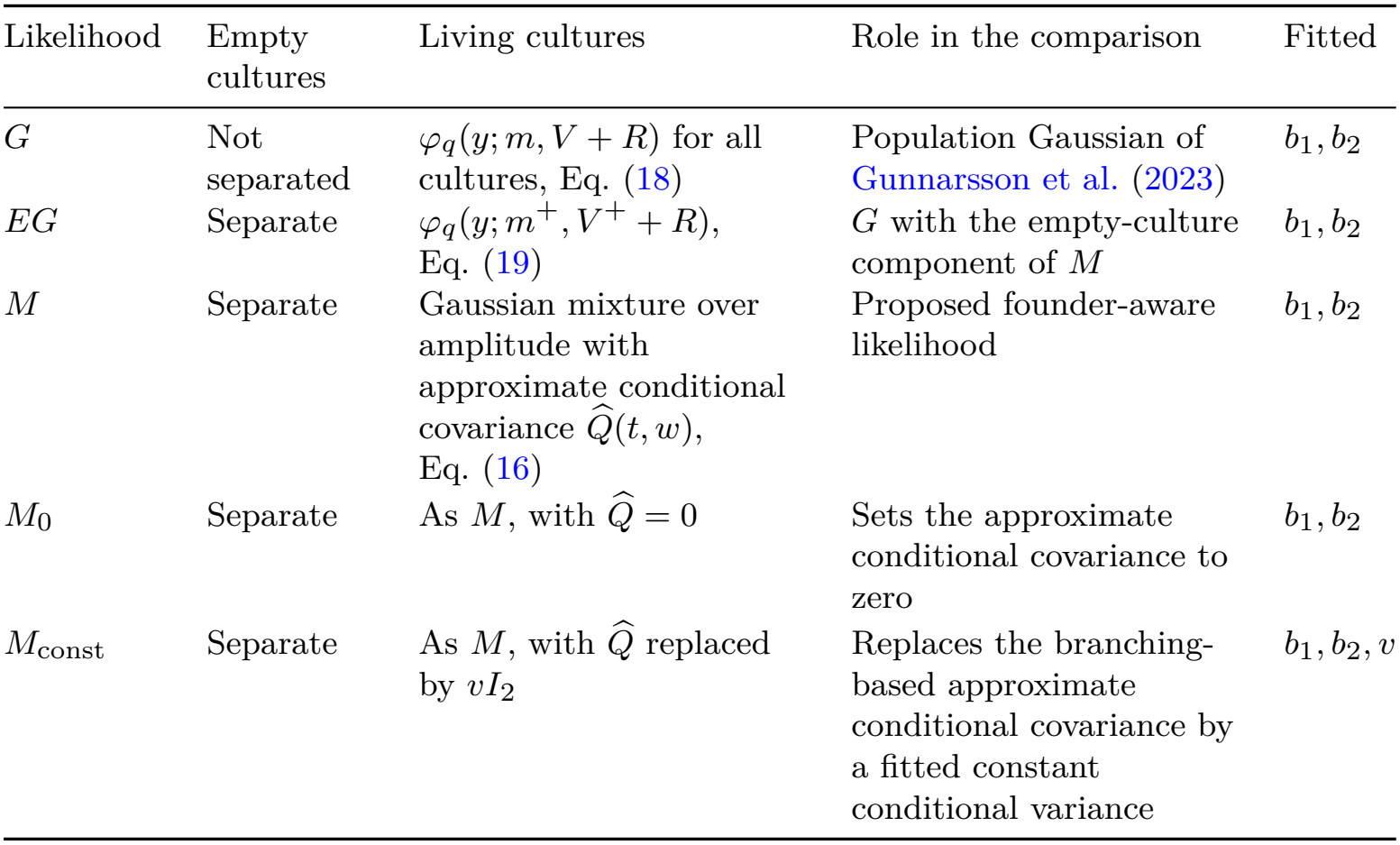
Likelihoods compared. Each observation density is evaluated for every observation and multiplied across cultures. All five likelihoods use the same observations, known inputs, measurement covariance *R*, and fitting region (Section 3.1). For the four likelihoods with a separate empty-culture component, this component is (1 *−π*_*t*_)*k*_0_(*y*), with *k*_0_(*y*) = *φ*_*q*_(*y*; 0, *R*), and the living-culture component has weight *π*_*t*_. The column “Role in the comparison” states what each comparator changes relative to *M*.

**Table 2.**
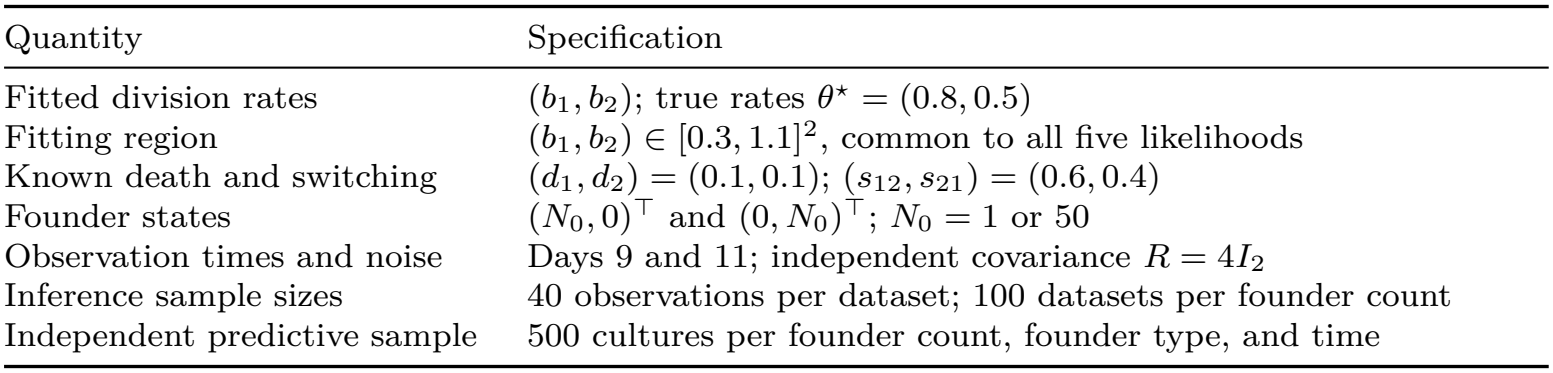
Main two-type birth–death experiment. Rates are per cell per day. Each dataset contains cultures with the same founder count, and results are reported separately for each founder count. All five likelihoods are fitted using the same observations and known inputs.

| Quantity | Specification |
|---|---|
| Fitted division rates | (b<sub>1</sub>, b<sub>2</sub>); true rates θ<sup>⋆</sup> = (0.8, 0.5) |
| Fitting region | (b<sub>1</sub>, b<sub>2</sub>) ∈ [0.3, 1.1]<sup>2</sup>, common to all five likelihoods |
| Known death and switching | (d<sub>1</sub>, d<sub>2</sub>) = (0.1, 0.1); (s<sub>12</sub>, s<sub>21</sub>) = (0.6, 0.4) |
| Founder states | (N<sub>0</sub>, 0)<sup>⊤</sup> and (0, N<sub>0</sub>)<sup>⊤</sup>; N<sub>0</sub> = 1 or 50 |
| Observation times and noise | Days 9 and 11; independent covariance R = 4I<sub>2</sub> |
| Inference sample sizes | 40 observations per dataset; 100 datasets per founder count |
| Independent predictive sample | 500 cultures per founder count, founder type, and time |

## 2 A founder-aware likelihood for endpoint observations

### 2.1 Establishment variation and the residual in the Yule process

We use an exact example with a single cell type to illustrate the distinction between variation in the establishment amplitude and variation in the residual conditional on that amplitude. Let *N* (*t*) be a Yule process (a pure-birth process) started from one cell, in which each cell divides at rate *b >* 0. Its expected size is E*N* (*t*) = exp(*bt*), and the population size divided by this growth factor converges almost surely to a random limit,

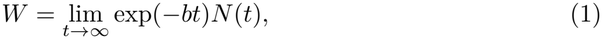

which has the exponential distribution with mean one. We call *W* the establishment amplitude, or the amplitude for short. Because *N* (*t*) is close to *W* exp(*bt*) at late times, the amplitude records the lasting effect of the earliest division times on population size. Conditional on the amplitude, the population has an exact representation due to Kendall (1966), stated as Theorem 2 in Athreya and Ney (1972, Chapter III, Section 11):

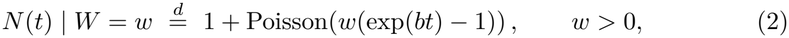

where 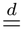 denotes equality in distribution and lowercase *w* is a specified value of *W*.

The amplitude determines the leading exponential growth, but the population still fluctuates around it. We define the residual at time *t* as the deviation from the leading exponential trajectory,

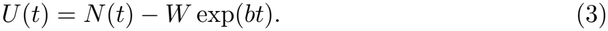

This definition does not condition on the amplitude, and the residual need not have zero conditional mean. Equation (2) gives the conditional mean and variance

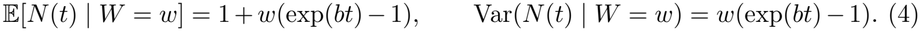

Consequently, E[*U* (*t*) |*W* = *w*] = 1 *−w* and Var(*U* (*t*) | *W* = *w*) = *w*(exp(*bt*) *−* 1). Subtracting *w* exp(*bt*) changes the conditional mean but leaves the conditional variance unchanged. Because *W* has mean and variance one, the law of total variance divides the population variance into variation of the conditional mean across amplitudes and the mean conditional variance:

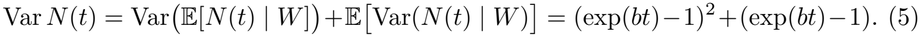

The first term, of order exp(2*bt*), comes from the distribution of the amplitude and dominates as the population grows. The second term, of order exp(*bt*), is the mean conditional variance of the residual given the amplitude. At late times, a Gaussian approximation with the population mean and variance must therefore mainly represent the amplitude distribution, which for one founder is exponential and strongly skewed. With *n* founders, the amplitude is the sum of *n* independent exponential amplitudes and has the Gamma distribution with shape *n* and scale one. Its skewness, 2*n*^*−*1*/*2^, decreases as the founder count increases, and by the central limit theorem the amplitude distribution approaches a Gaussian distribution.

Equation (5) motivates the construction in the rest of this section: use a Gaussian approximation to the conditional distribution of the residual given the amplitude, add back the leading exponential trajectory, and integrate over the distribution of the amplitude. The exact representation in Equation (2) serves as a benchmark for this approximation in Sections 2.3 and 4.

### 2.2 Multitype branching processes and observations

Let *Z*(*t*) = (*Z*_1_(*t*), …, *Z*_*q*_(*t*))^*⊤*^ be the random counts of *q* cell types at time *t ≥*0, with transpose denoted by *⊤*. We write *N* (*t*) for the total count, the sum of the entries of *Z*(*t*). The initial state *z*_0_ is known and nonzero, and the founder count is *N*_0_ = *N* (0). Each type-*j* cell evolves independently of all others and can undergo finitely many kinds of events. An event of kind *k* occurs at constant rate *α*_*jk*_ per cell and changes the vector of counts by a bounded integer vector *ν*_*jk*_. Division, death, and switching are events of this form. With *e*_*j*_ the *j*th coordinate vector, the mean-growth matrix *A* and the infinitesimal covariance at state *z* = (*z*_1_, …, *z*_*q*_)^*⊤*^ are

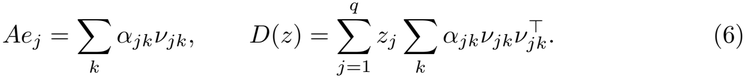

Write *m*(*t*) = E*Z*(*t*) = exp(*At*)*z*_0_ and *V* (*t*) = Cov(*Z*(*t*)) for the exact mean and covariance. The exact covariance satisfies *V* ^*′*^ = *AV* + *V A*^*⊤*^ + *D*(*m*) with *V* (0) = 0, where the prime denotes differentiation with respect to time *t*. The Yule process of Section 2.1 is the case *q* = 1 with division as the only event, so that *Z*(*t*) = *N* (*t*), *A* = *b*, and *D*(*z*) = *bz*.

Our test case is the two-type model of proliferation and reversible phenotypic switching studied by Gunnarsson et al. (2023). A type-*i* cell divides into two cells of its own type at rate *b*_*i*_, dies at rate *d*_*i*_, and switches to type *j* at rate *s*_*ij*_, for *i, j ∈ {*1, 2*}* with *i≠ j*. Rates are measured per cell per day, the initial state is *z*_0_ = (*n*_1_, *n*_2_)^*⊤*^, and *N*_0_ = *n*_1_ + *n*_2_. Equation (6) gives

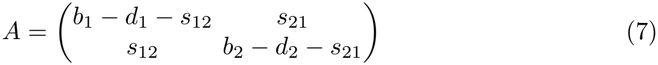

and

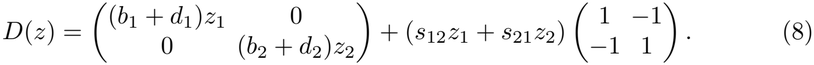

Each independently founded culture is observed once, at its assigned time, and we call the resulting vector *Y* the observation of that culture:

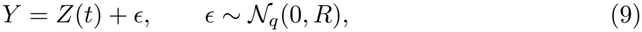

where *N*_*q*_ denotes a *q*-dimensional Gaussian distribution. Errors are independent of populations and across cultures (Gunnarsson et al. 2023, Equation (13)). In the two-type experiments, *R* = 4*I*_2_, where *I*_2_ is the identity matrix, so the standard deviation is two cells per component. This idealized measurement model applies to both living and empty cultures. Negative observations are retained without rounding, censoring, or classification. Survival status and cell counts are latent; inference uses only the observations, observation times, known founder states, and known model inputs. If the observation is a linear function of the counts, such as the counts of selected types or the total count, only the means and covariances of the Gaussian densities in the likelihood below need to change. Nonlinear functions of the counts, such as cell fractions, would require an additional approximation step.

### 2.3 Hypotheses, establishment amplitude, and conditional residual approximation

The construction uses four hypotheses.

(H1) Cells evolve independently, with finitely many kinds of events at constant finite rates and bounded changes in counts, as specified in Section 2.2.

(H2) The matrix *A* is irreducible, meaning that every type can eventually produce every other type.

(H3) The process is supercritical: the dominant eigenvalue *L* of *A* is positive. H4) Every other eigenvalue *λ* of *A* satisfies Re *λ < L/*2.

Under (H2), *A* has a real eigenvalue *L* that exceeds the real part of every other eigenvalue, with positive right and left eigenvectors *r* and *l*. We normalize these eigenvectors by *r*_*q*_ = 1 and *l*^*⊤*^*r* = 1. Under (H1)–(H3), the establishment amplitude

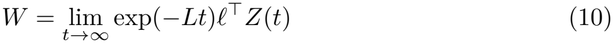

exists almost surely and in mean square (Athreya and Ney 1972; Janson 2004). The vector limit exp(*−Lt*)*Z*(*t*) *→ Wr* also holds almost surely. This almost sure limit is the continuous-time form of the theorem of Kesten and Stigum (1966b), stated by Georgii and Baake (2003, Theorem 2.1(b)). For *q* = 1 without death, Equation (10) reduces to Equation (1). The normalization of *l* sets the scale of *W* : its mean is *a*_0_ = E*W* = *l*^*⊤*^*z*_0_. Eventual survival, denoted by *S*, agrees with {*W >* 0} up to null events. The amplitude *W* is an unobserved random variable and is integrated out rather than estimated separately for each culture.

Extending Equation (3), we define the residual at time *t* by

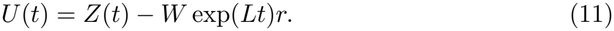

The family {*U* (*t*) : *t≥* 0} is the residual process. The definition is pathwise; conditioning is specified separately when discussing its distribution or moments.

Let *I*_*q*_ denote the *q× q* identity matrix, and define *P* = *rl*^*⊤*^ and *S* = *I*_*q*_ *−P*, the projections onto the dominant direction *r* and onto the complementary directions.

Conditional on *W* = *w >* 0, we approximate *Z*(*t*) by a Gaussian distribution with mean 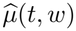 and covariance 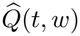, where

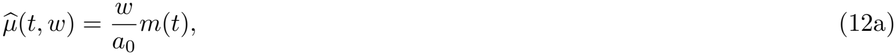

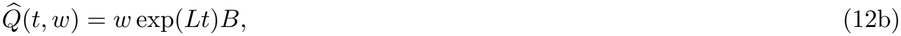

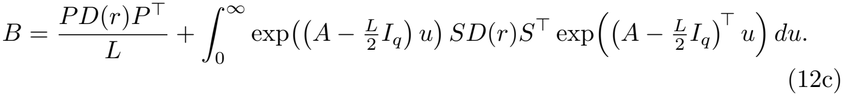

Since *Pz*_0_ = *a*_0_*r*, the exact population mean satisfies *m*(*t*) = *a*_0_ exp(*Lt*)*r* +exp(*At*)*Sz*_0_. Thus the centering 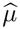 adds a finite-time adjustment to the leading exponential trajectory:

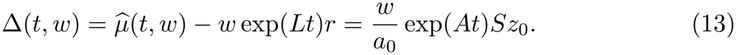

We call the Gaussian approximation to the conditional distribution of *U* (*t*) given *W* = *w >* 0 the conditional residual approximation. In Equation (12), the conditional distribution of *U* (*t*) given *W* = *w* is approximated by a Gaussian distribution with mean Δ(*t, w*) and covariance 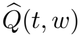. Adding back *w* exp(*Lt*)*r* gives the stated approximation to the population distribution. We call 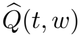 the approximate conditional covariance. It approximates the covariance of *U* (*t*) given *W* = *w*, which equals that of *Z*(*t*) because *w* exp(*Lt*)*r* is fixed once *W* = *w* is specified. The adjustment Δ retains the effect of the initial composition, but we do not assert that it equals the exact conditional mean of the residual.

The two terms of *B* in Equation (12c) have a direct interpretation. The first term accounts for events after time *t*, which determine the difference between *W* and the current scaled count exp(*−Lt*)*l*^*⊤*^*Z*(*t*); thus specifying *W* = *w* leaves *l*^*⊤*^*Z*(*t*) uncertain.

The second term accounts for demographic noise accumulated up to time *t* in the complementary directions, which produces random deviations of the composition of the population from the stable type proportions given by *r*. Section S1.2 of Online Resource 1 derives both terms from an exact decomposition of *U* (*t*). For any number of types, *B* is also the unique solution of the Lyapunov equation *FB* + *BF*^*⊤*^ + *D*(*r*) = 0, where 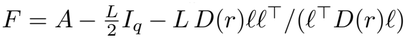. Thus, *B* can be computed by solving one linear system (Online Resource 1, Section S1.3).

The following theorem gives the joint limit of the amplitude and the scaled residual conditional on eventual survival as the observation time grows with the initial state fixed, which is limit (iii) in Section 1.

#### Theorem 1

*Assume (H1)–(H4) and fix a nonzero initial state z*_0_. *Let ξ be a q-dimensional Gaussian random vector with mean zero and covariance B, independent of W*. *Then, conditional on eventual survival W >* 0, *as t → ∞*,

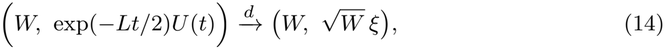

*where* 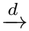 *denotes convergence in distribution of the pair under the law conditioned on W >* 0. *The same limit holds with U* (*t*) *replaced by* 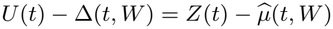.

Theorem 1 is an established result, which we state in our notation without proof: the limits in the complementary and dominant directions go back to Kesten and Stigum (1966a) and Athreya (1968, 1969), and the joint limit was stated by Asmussen and Hering (1983) and proved by Ren et al. (2014, 2017) and Kolesko et al. (2026). Section S1.2 of Online Resource 1 details the scope of each of these results.

Rescaling Equation (14) gives the approximation used in the living component of the likelihood. At late times, we approximate the distribution of *Z*(*t*) conditional on *W >* 0 by that of 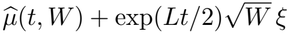, with *W* drawn from its law conditional on *W >* 0. This distribution is a mixture over the positive amplitude of Gaussian distributions with mean 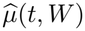 and covariance 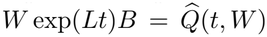. Because this mixture integrates over *W*, it uses only the joint limit conditional on eventual survival in Theorem 1 and requires no limit conditional on each specified value *W* = *w*. The theorem establishes convergence in distribution. It does not establish convergence of exact conditional moments, accuracy of densities or log likelihoods, or a bound on the error at finite times. Gaussian measurement noise smooths the count distribution, but noise with fixed covariance *R* vanishes on the theorem’s fluctuation scale after multiplication by exp(*−Lt/*2). Smoothing alone therefore does not establish the accuracy of the approximate observation density. We assess finite-time accuracy and the resulting inference empirically in Sections 3 and 4.

Hypotheses (H1)–(H3) give the amplitude, its exact moments, and the relation between eventual survival and a positive amplitude. Hypothesis (H4) is used for three specific purposes. First, it makes the integral in Equation (12c) finite and the Lyapunov equation for *B* uniquely solvable. Second, it makes the complementary component *SU* (*t*) of the residual of order exp(*Lt/*2), the square root of the population size, like the dominant component *PU* (*t*), so that both enter the Gaussian limit in Theorem 1. If instead Re *λ > L/*2 for some eigenvalue *λ*, the corresponding component grows like exp(Re *λt*) and has a random limit that is typically not Gaussian (Janson 2004, Theorem 3.1), so Equation (12b) would have the wrong scale. Third, it makes the adjustment Δ(*t, w*) in Equation (13) negligible on the scale exp(*Lt/*2) (Online Resource 1, Section S1.2). The population moments, survival probabilities, amplitude moments, and the comparison likelihoods *G* and *EG* of Section 2.5 do not require (H4). In the two-type model, positive switching rates imply (H2) and make the other eigenvalue *λ*_2_ of *A* real and distinct from *L*, because the discriminant (*A*_11_ *−A*_22_)^2^ + 4*s*_12_*s*_21_ of its characteristic polynomial is positive. Hypothesis (H4) then becomes *λ*_2_ *< L/*2, and *B* reduces to *PD*(*r*)*P*^*⊤*^*/L* + *SD*(*r*)*S*^*⊤*^*/*(*L −*2*λ*_2_) (Online Resource 1,

Section S1.3). When switching is much slower than net growth, *L* and *λ*_2_ are close to the larger and the smaller of the net growth rates *b*_1_ *−d*_1_ and *b*_2_ *−d*_2_, so (H4) fails whenever the slower type grows at more than about half the net rate of the faster type. Hypotheses (H3) and (H4) depend on the rates and are verified over the fitting region in Section 3.1.

For the Yule process (*q* = 1, *L* = *b, r* = *l* = 1, *a*_0_ = 1, *D*(*r*) = *b*, and *S* = 0), Equation (12) gives *B* = 1, so the approximate conditional mean and variance of the population are both *w* exp(*bt*). Here Δ(*t, w*) = 0, whereas the exact conditional mean of *U* (*t*) is 1*− w*. Subtracting the exact conditional mean and variance (Equation (4)) from their approximations gives *w−* 1 and *w*, respectively. These errors persist in absolute units even though the relative variance error, (exp(*bt*) *−* 1)^*−*1^, decreases with time. Section 4 examines the resulting quantile errors at early and late times.

Let *π*_*∞*_ = Pr(*S*) and *v*_0_ = Var(*W*) for the initial state *z*_0_, that is, for all *N*_0_ founders together. The amplitudes of independent founders add, so the moments for one founder give exact values of *a*_0_ and *v*_0_ for any initial composition (Online Resource 1, Section S1.4). Because *W* = 0 on extinction, the exact positive-amplitude moments are

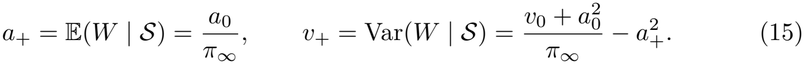

We approximate this distribution of the positive amplitude by a Gamma density 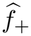 with shape 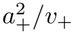 and scale *v*_+_*/a*_+_. The moments and the parameters of the Gamma distribution are recomputed for every candidate rate vector and initial state. The moment matching uses the total amplitude of the initial population, conditioned to be positive; it does not condition every founder to survive. The Gamma distribution is exact for the amplitude of the Yule process, for any founder count (Section 2.1), but is an approximation for the two-type population.

### 2.4 A likelihood retaining all seeded cultures

Let *A*_*t*_ = {*Z*(*t*) ≠ 0} denote finite-time survival and *π*_*t*_ = Pr(*A*_*t*_). These probabilities are calculated from the candidate branching parameters using backward extinction equations (Athreya and Ney 1972, Chapter V, Section 7.1) (Online Resource 1, Section S2.2). Let *φ*_*q*_(*y*; *µ, C*) denote a *q*-dimensional Gaussian density at the observed value *y*, with mean *µ* and covariance *C*, and write *k*_0_(*y*) = *φ*_*q*_(*y*; 0, *R*) for an empty culture’s observation density. The founder-aware likelihood, denoted by *M*, uses the observation density

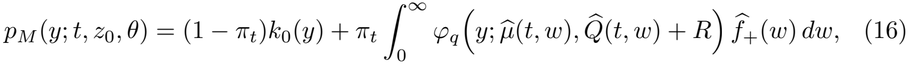

where *θ* comprises the fitted rates. In our experiments, *θ* = (*b*_1_, *b*_2_), and death and switching rates are known.

Integrating over positive amplitudes leaves conditioning on eventual survival within the living component. Using this component for finite-time survivors is a separate approximation, quantified by

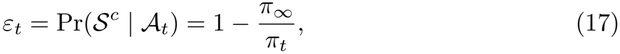

where *S*^*c*^ is eventual extinction. Holding the empty component fixed, replacing finite-time survivors by eventual survivors contributes at most *π*_*t*_*ε*_*t*_ in total variation distance between laws for all seeded cultures. This bound does not control relative density or log-likelihood error, nor the additional Gamma and Gaussian approximation errors. The weight of the living-culture component in Equation (16) is *π*_*t*_, not *π* _*∞*._

Each culture has its own independent amplitude. The integral over each culture’s amplitude is evaluated before densities are multiplied across cultures. Inside the integral, the Gaussian covariance is 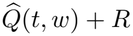, the approximate covariance of the residual conditional on *W* = *w* plus the measurement covariance. Variation between amplitudes is represented by the integral over *w* itself. Replacing 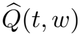 by the population covariance *V* (*t*), which already contains this variation (compare the first term of Equation (5)), would count amplitude variability twice.

### 2.5 Comparisons that separate extinction and residual effects

Table 1 summarizes the five likelihoods compared in this study.

The population Gaussian likelihood, denoted by *G*, uses the observation density

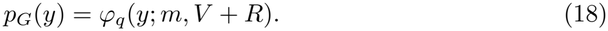

This density follows the likelihood of Gunnarsson et al. (2023), which uses the population mean and covariance. A calculation using their covariance formula agrees with our implementation (Online Resource 1, Section S4.2). The experiment is newly simulated and is not a reproduction of their published data analysis. As an estimation criterion, *G* is also a Gaussian pseudo-likelihood in the sense of Chen and Hyrien (2011, Section 3.3). Its score, the gradient *∇*_*θ*_ log *p*_*G*_(*y*) of the log observation density with respect to the fitted parameters *θ*, depends on the observation through linear and quadratic terms in *y− m*(*t*). Because *m* and *V* + *R* are the exact mean and covariance of the observation, the expected score at the true parameters is zero even when the observation distribution is non-Gaussian. With identifiability and suitable regularity conditions, this zero expected score supports consistent estimation even when the Gaussian density is inaccurate. Section 3.2 addresses calibration of the associated profile-likelihood confidence sets.

To distinguish mixing over the amplitude from explicit representation of empty cultures, we also use *EG*, an adaptation of *G* with exactly the same empty-culture component as the founder-aware likelihood *M* in Equation (16):

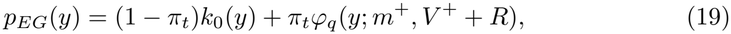

where *m*^+^ = *m/π*_*t*_ and *V* ^+^ = (*V* +*mm*^⊤^)*/π*_*t*_ *−m*^+^*m*^+⊤^ are exact moments conditional on *A*_*t*_. Both *G* and *EG* recover the exact population mean and covariance before adding measurement noise. With no death, *π*_*t*_ = 1 and *G* and *EG* coincide.

For the mixture likelihoods *EG* and *M*, agreement of overall moments alone does not give the zero expected score established for *G*.

Two variants of the founder-aware likelihood retain its amplitude distribution, approximate conditional population mean 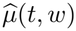, and empty-culture component. The variant *M*_0_ sets the approximate conditional covariance 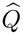 to zero. In its living component, the population is fixed at 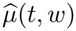 given *W* = *w*, so the residual is fixed at Δ(*t, w*); measurement error remains in the observation model. The variant *M*_const_ replaces 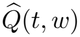 by *vI*_2_ and fits *v ≥* 0 jointly with the birth rates. This single variance is shared across founder types and observation times within a dataset. The variance *v* is a fitted constant conditional variance for living cultures; the known measurement covariance remains *R* for both living and empty cultures. Thus *M*_const_ has three fitted parameters, while the other likelihoods have two, and *v* = 0 recovers *M*_0_. Under *M*_const_, the counts of the two types in a culture still depend on one another because they share a latent amplitude.

The comparison with *M*_0_ tests the effect of retaining 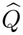. By comparing *M* with *M*_const_, we assess what the approximate conditional covariance derived from the branching model adds beyond a fitted constant conditional variance. Replacing 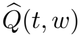 by *vI*_2_ changes amplitude dependence, time dependence, direction, and correlation together, so this comparison does not isolate any one feature of the covariance.

The variant *M*_0_ also connects our comparison to an existing estimation method. For the one-founder Yule process without measurement error, *M*_0_ places the population at 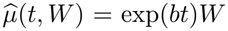, which has the exponential density with mean exp(*bt*). The logarithm of this density is the exponential pseudo-likelihood of Chen and Hyrien (2011, Section 3.3, Eq. (7), and Section 4.1), who used this density for estimation without assuming that it is the exact distribution of the data. More generally, *M*_0_ has the structure of likelihoods that integrate over a random amplitude or time shift, conditional on which the trajectory is deterministic. Morris et al. (2025, Equations (6) and (7)) use such a likelihood with a random time shift, for populations that do not go extinct, with independent Gaussian observation errors on the logarithmic scale. The comparison of *M* with *M*_0_ therefore examines what the approximate conditional covariance 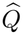 adds to these approximations.

## 3 Numerical assessment

### 3.1 Design and reference calculations

Table 2 specifies the main experiment, which we call the birth–death experiment to distinguish it from the experiments without death described later in this section. At each founder count, each inference dataset contains 40 independent endpoint observations: ten cultures for each combination of two pure founder types and two times. We call the birth rates used for simulation the true rates, 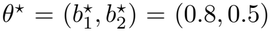 per day.

We generate 100 datasets per founder count at the true rates with the stochastic simulation algorithm of Gillespie (1977), which simulates every division, death, and switching event exactly, and add to each culture an independent measurement error drawn from Equation (9). Every culture is retained, and all five likelihoods are fitted to the same datasets. The true rates are used to simulate data and to compute estimation errors and coverage. Neither the true rates nor the simulated cell counts and survival status are available to fitting. All estimates and profile-likelihood confidence sets are computed within the same fitting region, the set of birth-rate pairs (*b*_1_, *b*_2_) *∈* [0.3, 1.1]^2^, with death and switching rates fixed at their known values. Section 5 discusses the choice of known rates.

For every rate vector in the fitting region, with the known death and switching rates, *A* is irreducible, *L >* 0, and *λ*_2_ *< L/*2, so hypotheses (H2)–(H4) hold throughout the region (Online Resource 1, Section S2.1). The probability *ε*_*t*_ in Equation (17), which measures the effect of using eventual survivors in place of finite-time survivors, is small at the true and fitted rates but larger near the lower boundary of the fitting region. Section S2.3 of Online Resource 1 gives these values.

An independent predictive sample compares the observation densities of *G, EG, M*, and *M*_0_ at the true rates. For each observation, we calculate the log-density difference log *p*_*M*_ (*Y*) *−*log *p*_*H*_ (*Y*) for comparator *H ∈ {G, EG, M*_0_ }. Positive mean differences favor *M*. We calculate standard errors from paired differences on the same cultures. The predictive comparison evaluates distributions at the true rates; it is not prediction after parameter fitting. Because *M*_const_ requires a variance fitted to data, it is not included in the predictive comparison.

Two experiments without death (*d*_1_ = *d*_2_ = 0) serve as controls in which no culture becomes empty, so an advantage cannot come from the treatment of empty cultures. The two experiments differ only in their observation days, days 5 and 7 in one and days 9 and 11 in the other. We refer to each pair of observation days as a window. Otherwise, both use the same true birth rates, switching rates, measurement covariance, founder states, and inference sample sizes as the birth–death experiment, and each has its own independently simulated datasets. The window of days 9 and 11 matches the birth– death experiment. We report both windows in order to avoid selecting observation times that favor one likelihood. The two experiments do not isolate the causal effect of observation time. These experiments compare *G, M*, and *M*_0_. Without death, *EG* coincides with *G*, and *M*_const_ was fitted only in the birth–death experiment.

Exact reference calculations use the Yule process of Section 2.1 with *b* = 0.8, one founder, and times 0.5 and 5 days. Because this process has no death and its amplitude distribution is exact, the comparison isolates error in the conditional residual approximation. We compare each approximate distribution with the exact distribution through quantiles. For each likelihood, we plot the quantiles of its approximate distribution against those of the exact distribution at 13 probability levels from 0.001 to 0.999. Agreement places the points on the diagonal. We summarize each comparison by the maximum absolute difference between approximate and exact quantiles over these levels. The comparison is made for the cell count *N* (*t*), giving cell-count quantiles, and for the observation *Y* = *N* (*t*) + *ϵ* with *ϵ∼N* (0, 4), giving observation quantiles. For cell-count quantiles, each approximate distribution is its observation density with the measurement variance set to zero, so its Gaussian components can produce negative quantiles. Section S3 of Online Resource 1 gives the exact distribution and the approximate distributions compared. Quantile agreement alone establishes neither density accuracy nor the ranking of likelihoods in parameter estimation, so we assess these outcomes separately.

### 3.2 Estimation, uncertainty, and numerical accuracy

For each of the five likelihoods, we estimate the birth rates by maximizing the sum of log observation densities over the fitting region. Each maximization starts from several points of a grid over the region, uses bounded optimization, and checks the boundary of the region explicitly. For *M*_const_, we maximize the log likelihood jointly over the variance *v≥* 0 and the birth rates. An upper bound on the likelihood shows that sufficiently large values of *v* cannot attain the maximum, so no upper limit on *v* is imposed in advance (Online Resource 1, Section S4.1).

Let ℒ_*H*_ be the log likelihood of a dataset under *H ∈ {G, EG, M, M*_0_, *M*_const_ }, and let 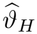 be the parameter vector that maximizes it. Also let 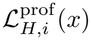 be the maximum of ℒ_*H*_ with *b*_*i*_ = *x* fixed and all other fitted parameters reoptimized: the other birth rate, and also *v* for *M*_const_. The nominal 90% profile-likelihood confidence set for *b*_*i*_, which we call the profile set for short, is

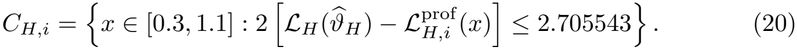

The cutoff is the 90th percentile of a chi-square distribution with one degree of freedom. Following the unadjusted chi-square profile rule used by Gunnarsson et al. (2023, Appendix A.2, Equation (20)), we apply this rule to every likelihood and assess its coverage empirically. If the likelihood is the true density of each observation, the variance of the score equals the expected curvature of the log likelihood. Under standard regularity conditions, this equality makes the chi-square cutoff valid asymptotically as the number of independent cultures grows. For Gaussian pseudo-likelihood, the asymptotic covariance of the estimates generally has a sandwich form, which accounts separately for the variability of the score and the curvature of the criterion (Chen and Hyrien 2011, Section 3.3). Because the score of a Gaussian density is quadratic in the observation, its variance depends on the third and fourth moments of the observation. Correct first and second moments of the observation therefore do not by themselves justify this chi-square cutoff for profile sets (Kent 1982). Our coverage comparison concerns the common unadjusted rule, not Gaussian inference with sandwich-adjusted uncertainty. We also assess coverage under *M* for the simulated endpoint experiments; empirical agreement with the nominal level does not establish general chi-square calibration. Coverage records whether the true rate belongs to the set. If a set consists of several intervals, its total length is the sum of their lengths.

We report bias, root-mean-square error (RMSE), coverage, and mean total length of the profile sets of the estimated birth rates. To compare *M* with a comparator *H*, we report three differences: the percentage reduction in RMSE, 100(1 *−*RMSE_*M*_ */*RMSE_*H*_), which is positive when *M* has the smaller RMSE; the coverage gain, coverage under *M* minus coverage under *H* in percentage points; and the percentage increase in mean total length from *H* to *M*. We call these differences paired because both likelihoods are fitted to the same datasets. Their uncertainty is estimated by a paired bootstrap that resamples whole datasets, so every resample compares the likelihoods on the same observations (Online Resource 1, Section S5). The resulting 95% intervals describe Monte Carlo uncertainty in differences between likelihoods and are distinct from the nominal 90% profile sets computed for each dataset. We call a difference resolved when its 95% interval excludes zero. Each interval is computed for one comparison and is not adjusted for multiple comparisons. With 100 datasets, coverage near 90% has a Monte Carlo standard error of about three percentage points.

The integral over amplitude in Equation (16) is computed by adaptive numerical integration in the variable log *w* (Piessens et al. 1983; Virtanen et al. 2020). Representative density evaluations, fits, and profiles are repeated with tighter tolerances, smaller omitted amplitude tails, and denser grids, including small and negative observed values and candidate rates near the boundary of the fitting region. Sample means and covariances of simulated cultures agree with the exact moments *m*(*t*) and *V* (*t*) within Monte Carlo error, so the simulator and the moment calculation check each other. Section S4 of Online Resource 1 reports this check, the numerical settings, and the measured changes. In the birth–death experiment, all 1,000 fits (five likelihoods for each of 200 datasets) and all 2,000 profile sets (two birth rates for each fit) were computed successfully. No dataset was excluded, and no profile set was truncated at the boundary of the fitting region. Failed optimization attempts from individual starting points are recorded even when the search as a whole succeeded.

## 4 Results

### 4.1 Exact Yule comparison

The exact Yule comparison shows why mixing over the amplitude can matter even when population size is large (Figure 1). At day 5, the approximate observation quantiles of *M* and *M*_0_ lie close to the exact quantiles, whereas those of the population Gaussian likelihood *G* deviate by up to about 150 cells (Figure 1D). *G* poorly represents the skewness inherited from a single founder. The approximate cell-count quantiles show the same pattern (Figure 1B): at day 5, the 0.001-quantile of *G* is about *−*113 cells, although a Yule population started from one cell always contains at least one cell. At day 0.5, however, the order is reversed, and the approximate observation quantiles of *G* are closer to the exact quantiles than those of *M* and *M*_0_ (Figure 1C). Adding measurement noise smooths the distribution but does not remove the error of the conditional Gaussian approximation at early times.

**Fig 1.**
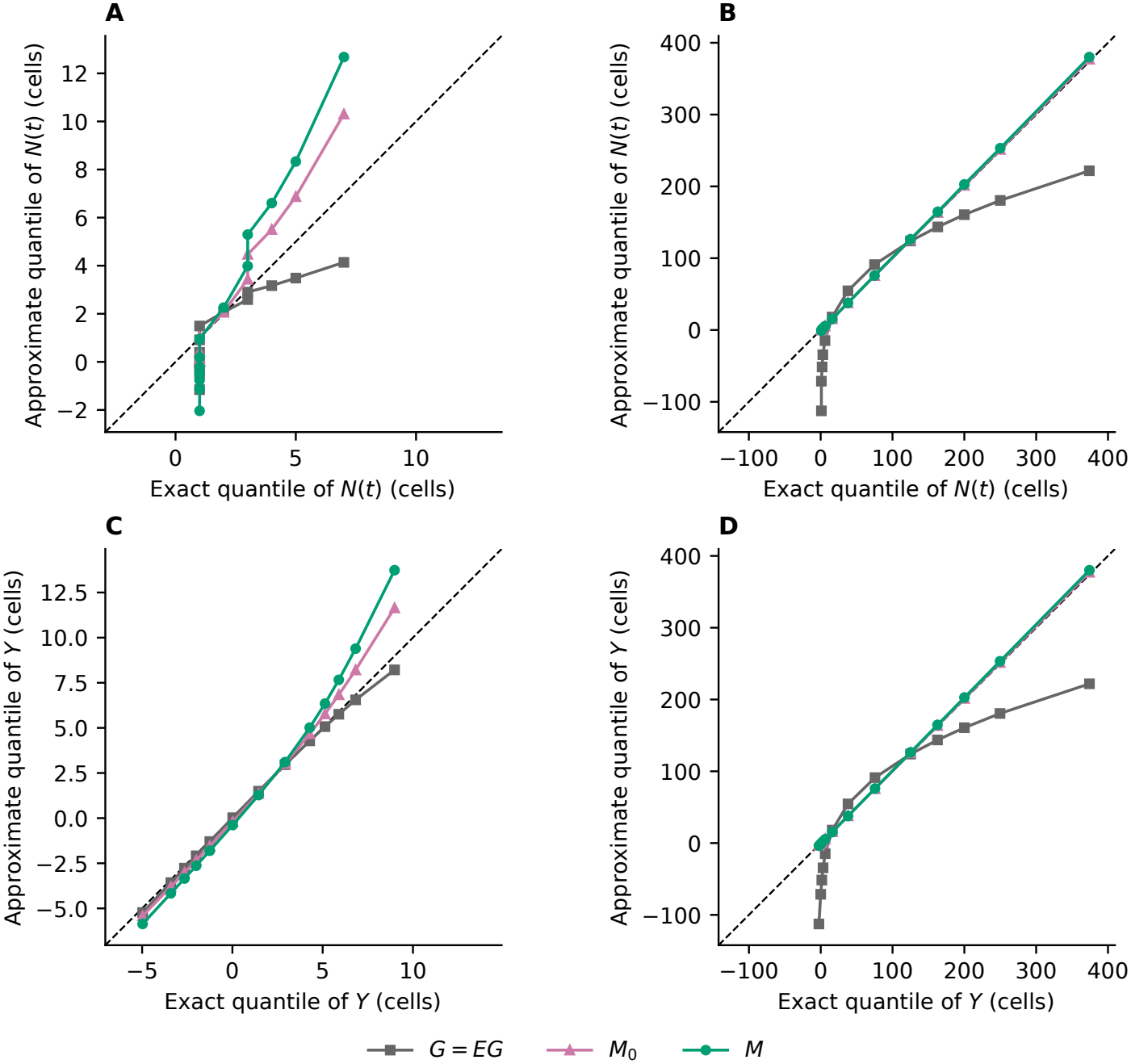
Quantile comparison for the one-founder Yule process at early and late times. Each curve plots quantiles of an approximate distribution against quantiles of the exact distribution at 13 probability levels from 0.001 to 0.999 (Section 3.1). The dashed diagonal denotes equality. (A) Cell count *N* (*t*) at day 0.5. (B) Cell count *N* (*t*) at day 5. (C) Observation *Y* = *N* (*t*) + *ϵ* at day 0.5. (D) Observation *Y* = *N* (*t*) + *ϵ* at day 5. In A and B, each approximate distribution is evaluated with the measurement variance set to zero, and negative approximate quantiles are retained; in C and D, *ϵ∼ N* (0, 4) independently of *N* (*t*). The division rate is *b* = 0.8 per day, and the amplitude distribution is exact in this example. The likelihoods are summarized in Table 1. Without death, *G* and *EG* coincide. *M*_const_ is not shown because the rates of the branching model alone do not specify its distribution; a value of its fitted constant conditional variance *v* is also required. Axes have equal scales within each panel.

At both times, the maximum absolute difference between approximate and exact observation quantiles is smaller for *M*_0_ than for *M* (Figure 1C, D).

### 4.2 Predictive gains beyond empty cultures

We first compare observation densities on the independent predictive sample of the birth–death experiment, evaluated at the true rates (Section 3.1). With one founder, *M* assigns higher mean log density to the observations than both *G* and *EG* in all four combinations of founder type and observation time (Figure 2A). The mean of log *p*_*M*_ (*Y*) *−*log *p*_*G*_(*Y*) per culture lies between 1.0 and 1.5, the mean of log *p*_*M*_ (*Y*) *−* log *p*_*EG*_(*Y*) is about half as large, and every mean difference is more than eight Monte

**Fig 2.**
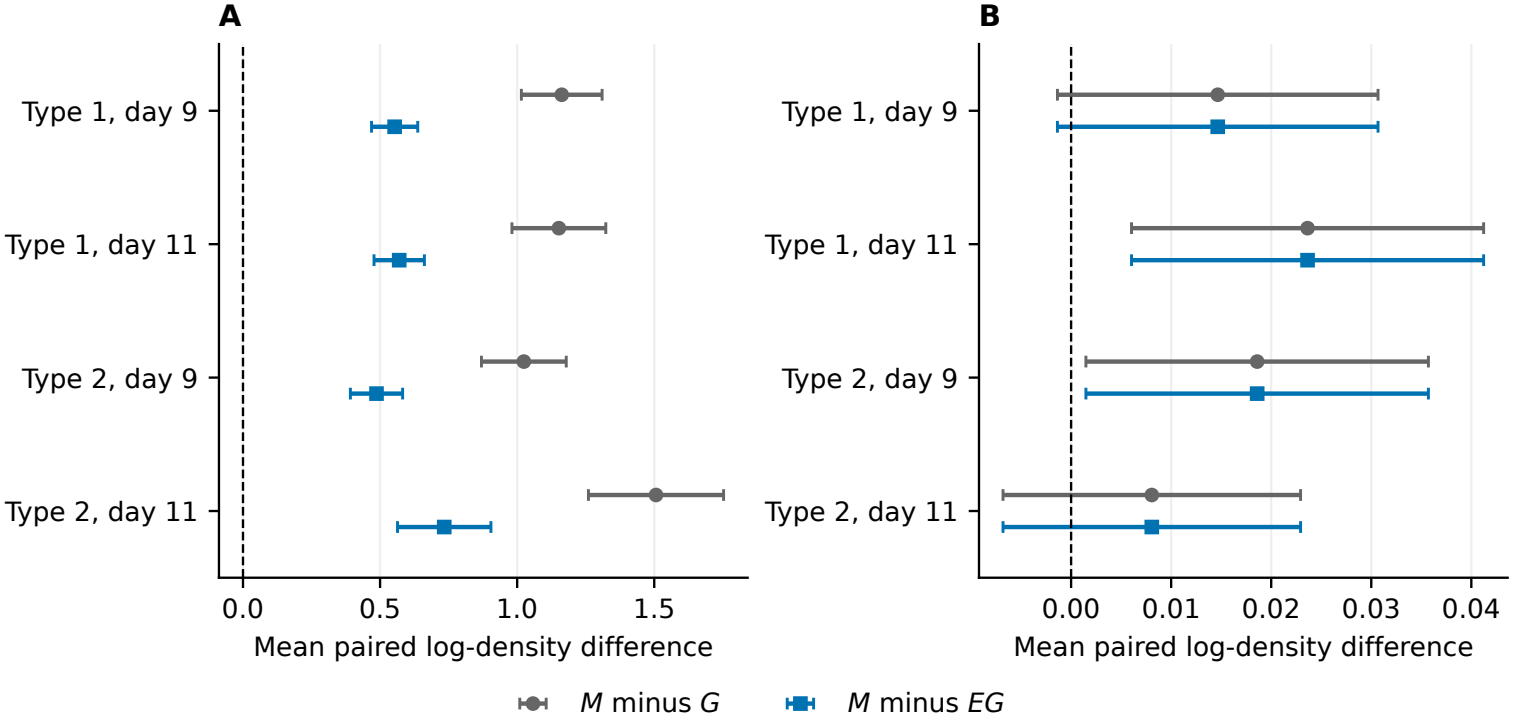
Paired differences in log observation density on the independent predictive sample of the birth– death experiment (Table 2), at the true rates. Each point is the mean difference over 500 independent observations with the indicated founder type and time. Bars are 1.96 Monte Carlo standard errors. Positive differences favor *M*. Differences from *M*_0_ are omitted because they are much larger (Online Resource 1, Table S4). (A) One founder per culture. (B) 50 founders per culture, with a different horizontal scale to resolve the much smaller differences. The comparisons with *G* and *EG* essentially coincide and are offset vertically for visibility. These densities are evaluated at the true rates, not at fitted rates.

Carlo standard errors from zero. Modeling empty cultures explicitly therefore explains only part of the gain of *M* over *G*. Because *M* and *EG* also differ in the mean and covariance they assign to living cultures, the remaining gain cannot be attributed exclusively to non-Gaussian shape.

With 50 founders, the mean differences of log *p*_*M*_ (*Y*) from log *p*_*G*_(*Y*) and from log *p*_*EG*_(*Y*) are below 0.03, and two of the four combinations are within 1.96 Monte Carlo standard errors of zero (Figure 2B). Relative to *M*_0_, which sets 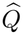 to zero, *M* has much larger gains in log density with both founder counts (Online Resource 1, Table S4).

### 4.3 Better estimation and coverage with one founder

Figure 3 shows the inference results of the birth–death experiment (Table 2) for both birth rates and founder counts. It reports the percentage reductions in RMSE under *M* relative to each comparator, together with the coverage and mean total length of the nominal 90% profile sets (Equation (20)) for all five likelihoods. Coverage is the percentage of the 100 datasets whose profile set contains the true rate (Section 3.2). Table S5 of Online Resource 1 gives the paired differences defined in Section 3.2, with their 95% intervals. With one founder per culture, *M* estimates the type-1 birth rate *b*_1_ more accurately than either Gaussian likelihood (Figure 3A). Its RMSE for *b*_1_ is 17% lower than under *G* (95% interval 10–23%), with a similar reduction relative to *EG*. If RMSE decreases in proportion to the inverse square root of the number of cultures, *G* would need about 1.4 times as many cultures to match the *b*_1_ RMSE of *M*. For *b*_2_, *M* also has the smallest RMSE among *G, EG*, and *M*, but the reductions are not resolved.

**Fig 3.**
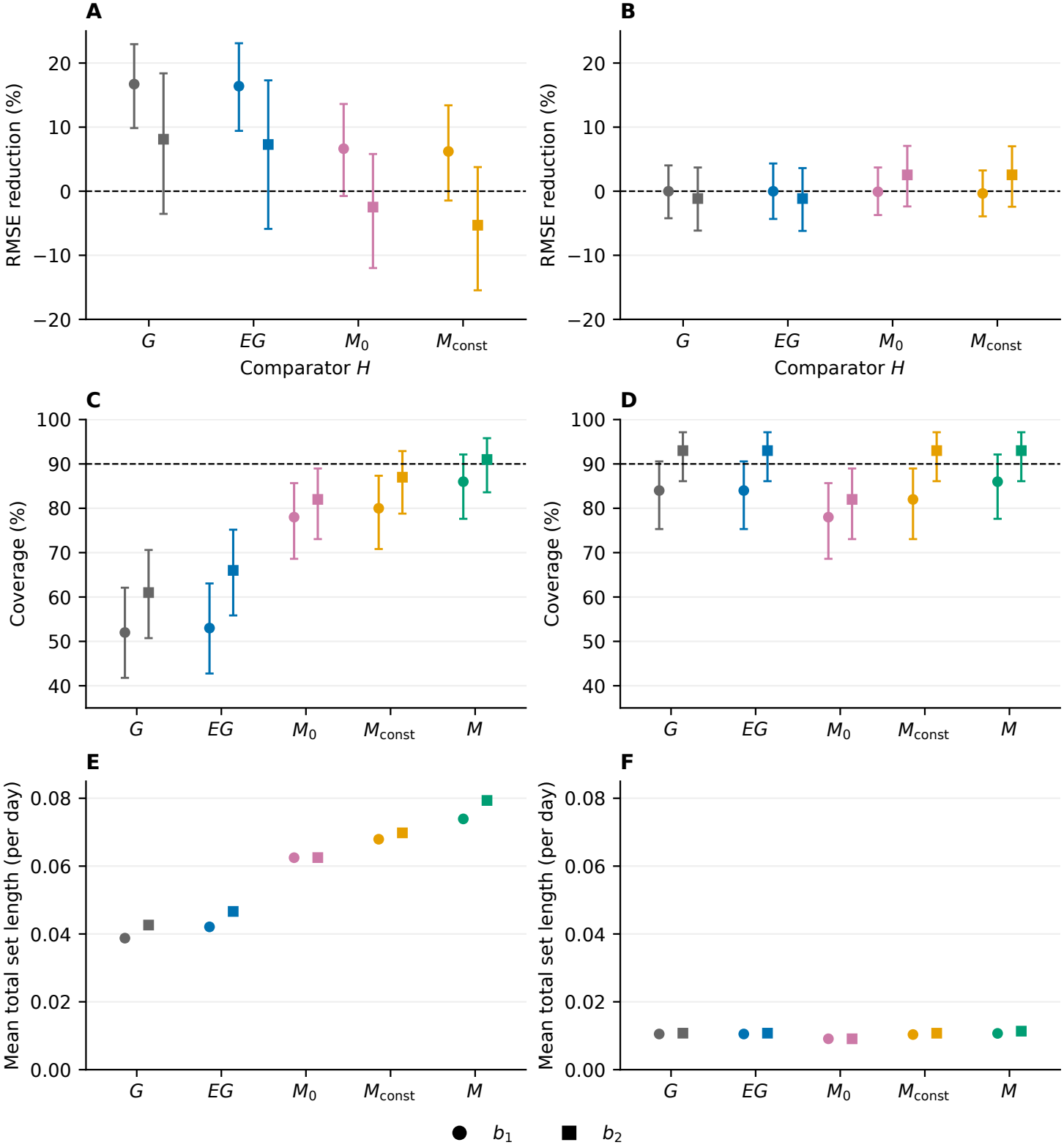
RMSE reductions, coverage, and length of profile sets for the five likelihoods in the birth– death experiment (Table 2). (A) RMSE reduction with one founder per culture. (B) RMSE reduction with 50 founders per culture. (C) Coverage of nominal 90% profile sets with one founder, under the common unadjusted chi-square rule. (D) Coverage with 50 founders. (E) Mean total set length with one founder. (F) Mean total set length with 50 founders. Circles and squares indicate the two fitted birth rates. Both columns use the same vertical scale within each row. RMSE reduction is 100(1*−* RMSE_*M*_ */*RMSE_*H*_) relative to comparator *H*; positive values favor *M*. RMSE bars are paired 95% bootstrap intervals over 100 datasets, coverage bars are exact 95% binomial intervals (Clopper and Pearson 1934), and length bars are one Monte Carlo standard error. Comparisons of coverage and length use paired differences, rather than overlap of these bars (Online Resource 1, Table S5). The dashed lines mark zero RMSE reduction and 90% coverage. Table S6 of Online Resource 1 gives absolute RMSEs with individual 95% bootstrap intervals, and signed biases.

With one founder per culture, the nominal 90% profile sets under *M* cover the true rate in 86% of datasets for *b*_1_ and 91% for *b*_2_, whereas the sets under *G* and *EG* cover it in only 52–66% (Figure 3C). The exact 95% binomial intervals for the coverage of *M* include the nominal level. The sets under *M* are about 1.9 times as long as those under *G* (Figure 3E). The gain is in calibration rather than precision: the sets of *G* are shorter but miss the true rate far more often than the nominal level allows.

With 50 founders per culture, the many-founder setting for which *G* was developed, the likelihoods give similar results (Figure 3B, D, F). No difference in RMSE or coverage between *M* and either Gaussian likelihood is resolved, and the sets under *M* are at most 6% longer. The founder-aware likelihood thus improves inference with one founder per culture and performs comparably with 50 founders.

### 4.4 Retaining the approximate conditional covariance improves coverage

In the birth–death experiment with one founder per culture, coverage is higher under *M* than under *M*_0_, which omits the approximate conditional covariance 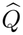. Coverage rises from 78% and 82% to 86% and 91% for *b*_1_ and *b*_2_, respectively (Figure 3C). Both coverage gains are resolved and come with longer sets. The RMSE differences between *M* and *M*_0_ are not resolved. At 50 founders, coverage under *M* also exceeds that under *M*_0_ (Figure 3D), while the RMSE differences remain unresolved.

*M*_const_, which uses a fitted constant conditional variance, recovers part of these coverage gains. With one founder, its coverage for *b*_1_ is 80%, versus 86% for *M*, a resolved gain of 6 percentage points for *M* with sets about 9% longer; for *b*_2_, the coverage difference is not resolved (Figure 3C). At 50 founders, *M* retains a smaller resolved coverage gain for *b*_1_, and the two likelihoods have equal coverage for *b*_2_ (Figure 3D).

No RMSE comparison between *M* and *M*_const_ is resolved (Online Resource 1, Table S5). The 100 datasets thus do not determine which of the two likelihoods estimates the birth rates more accurately, and they also do not show that the two are equally accurate. Retaining 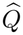 therefore improves coverage. The endpoint results do not establish whether the full approximate conditional covariance derived from the branching model is needed to achieve the RMSE observed under *M*. Section 5 discusses observation designs that could separate *M* from *M*_const_.

### 4.5 Experiments without death

In the two experiments without death (Section 3.1), no culture can be empty, and *M* again improves inference with one founder (Online Resource 1, Table S7). In both windows of observation days, *M* has a resolved RMSE reduction relative to *G* for at least one birth rate, and higher coverage with longer sets. These experiments also give the only resolved RMSE reductions of *M* relative to *M*_0_, about 12% and 19% for *b*_2_ at days 9 and 11 and at days 5 and 7, respectively.

### 4.6 Computational cost

For each candidate rate vector, *G* evaluates one Gaussian density per observation. The founder-aware likelihood *M* additionally requires finite-time and eventual extinction probabilities for each founder state and observation time, and a numerical integral over amplitude for each observation. Absolute times depend on hardware and implementation, so we compare likelihoods by ratios of times measured on the same saved datasets of the birth–death experiment with the same execution settings. Section S4.3 of Online Resource 1 gives the absolute times and settings. In a matched comparison of *G, EG, M*, and *M*_0_, a complete fit under *M* took 5.0–6.5 times as long as under *G*, with the range over the two founder counts. Table S3 of Online Resource 1 gives the corresponding times for single likelihood evaluations and profile calculations. Most of the additional fitting time was spent computing extinction probabilities rather than integrating over amplitude, and fits under *EG*, which requires the same extinction probabilities, took about as long as under *M*.

In a separate matched comparison, fits under *M*_const_ took about 5–7 times as long as under *M*, and profile calculations about 21–23 times as long, mainly because the additional variance *v* is reoptimized at every profile point. Fitting *M* and computing its profile sets can therefore cost less than the corresponding calculations for *M*_const_, despite the more detailed approximate conditional covariance of *M*.

## 5 Discussion

Two features of *M* account for its differences from *G* and from *M*_0_. First, with one founder, the variance of the endpoint population is dominated by variation of the establishment amplitude (Equation (5)), whose distribution is strongly skewed. The Gaussian density of *G* has the exact mean and covariance of the endpoint population, but it cannot represent the skewness that this amplitude variation produces in the population (Figure 1B, D). The narrow profile sets and low coverage under *G* are consistent with this misspecification. As the founder count *N*_0_ grows, the skewness of the amplitude decreases in proportion to 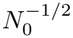 and the mixture approaches a single Gaussian distribution, consistent with the comparable performance of *M* and *G* at 50 founders.

Second, specifying the amplitude does not fix the endpoint population or its composition. Under *M*_0_, every component of the living part of the mixture has its mean on the ray {*w m*(*t*)*/a*_0_ : *w >* 0} and covariance *R*. Thus, the component of an observation perpendicular to *m*(*t*) has only the measurement variance, whatever the amplitude. In the branching process, however, demographic variation in that direction grows with population size. The one-type Yule process has no such perpendicular direction, which is consistent with the approximate observation quantiles of *M*_0_ matching the exact quantiles more closely than those of *M* (Figure 1C, D). In the two-type model, by contrast, *M* assigns much higher predictive log density than *M*_0_, with an advantage that grows roughly in proportion to the expected population size (Online Resource 1, Table S4). Retaining 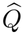 also widens the profile sets, which improves coverage (Section 4.4).

Although *M*_0_ assigns much lower predictive log density than *M*, the difference in their RMSEs is not resolved. Coverage under *M*_0_ is lower than under *M*, but it is much closer to that of *M* than to that of *G* (Figure 3A, C). A possible explanation, which we have not tested, is that the composition of a culture provides little information about the birth rates in this design. The estimates would then depend mainly on population sizes, whose distribution is dominated by the amplitude distribution that *M*_0_ and *M* share.

Replacing 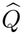 by a fitted constant conditional variance, instead of setting it to zero, recovers part of the coverage gain from the approximate conditional covariance derived from the branching model. Coverage under *M* remains higher than under *M*_const_ for *b*_1_, with a resolved difference and longer sets, but the RMSE difference is not resolved (Section 4.4). Each culture is observed at only one of the two times, so the data inform the conditional covariance at each time but not the covariance between longitudinal observations of one population. In this design, one constant variance may approximate the residual variation well enough to give similar rate estimates, although the unresolved RMSE comparison cannot rule out differences too small to detect with 100 datasets. A wider separation between observation times could expose time dependence that one constant variance cannot reproduce, but it would also change population sizes and approximation error, so a benefit is not guaranteed. The endpoint evidence therefore supports the founder-aware likelihood as a whole without establishing that each feature of its approximate conditional covariance is needed.

Our study examines how founder count affects likelihood performance, complementing the result of Chen and Hyrien (2011), which established consistency as the number of independent cultures increases. The population Gaussian likelihood *G* can support consistent estimation from increasing numbers of independent cultures without becoming an accurate density for each culture (Gourieroux et al. 1984; Chen and Hyrien 2011). This robustness follows from the exact moments of *G* and is a useful property. It does not follow automatically for *M*, which uses a Gamma approximation to the amplitude distribution and a conditional Gaussian approximation. Exact moments do not, however, justify the chi-square cutoff for *G* (Section 3.2). The variability of its estimates also depends on higher moments, which a Gaussian density misrepresents when the amplitude distribution is skewed, so its profile sets can be too narrow.

All coverage results use the common unadjusted profile rule of Section 3.2. We did not compare *M* with Gaussian pseudo-likelihood whose uncertainty is adjusted for distributional misspecification. We could assess whether the coverage gain of *M* persists by comparing it with *G* using sandwich-based uncertainty. That comparison would evaluate coverage together with interval length under the same observation design. Our comparison also does not include exact maximum likelihood, which Chen and Hyrien (2011, Section 4.2 and Table 2) found substantially more accurate than Gaussian pseudo-likelihood for single-founder birth–death populations observed longitudinally. The two-type process with reversible switching has no known closed-form likelihood, and evaluating it numerically at every candidate rate vector during fitting and profiling would be computationally expensive (Xu et al. 2015; Ho et al. 2018). Saddlepoint approximations to the distribution of population sizes in multitype branching processes (Luo et al. 2026) offer a numerical route to such a comparison, although their stated model changes type through births of a new type rather than through switching. We did not attempt this calculation, so we cannot determine how much *M* reduces the difference in accuracy between Gaussian pseudo-likelihood and exact maximum likelihood.

The comparators of Section 2.5 isolate the effect of each part of the founder-aware likelihood, because successive likelihoods in Table 1 differ in a single component. We also gave all five likelihoods the same known death and switching rates and fitted only the two birth rates, so that the comparison does not depend on whether the observation design identifies every rate. With death and switching rates known, the birth rates determine the net growth rate of each type, and Gunnarsson et al. (2023, Section 4.1 and Table 2) showed that net growth rates and switching rates are identifiable from mean cell counts. The net growth rates are also the parameters confounded with the establishment amplitude in a single endpoint observation, where faster growth and a favorable early history both produce a larger culture. The low coverage of the profile sets under *G* therefore occurs for parameters that are identifiable from mean counts. In practice, death and switching rates are usually unknown, and separating division from death requires information beyond the mean (Gunnarsson et al. 2023, Sections 4.1 and 4.2.3), so the approximation of the data distribution could matter more when all rates are fitted. Such a comparison would also depend on whether the observation design contains enough information to separate the rates (Gunnarsson et al. 2023, Section 4.2.3).

We used maximum likelihood because it compares the approximate likelihoods with *G* under common observations and the same fitting region, without introducing differences due to prior distributions. The same likelihoods could be used in Bayesian and hierarchical inference, as Morris et al. (2025) did with a random time shift, but posterior credible sets would then need their own assessment of repeated-sampling coverage.

The multitype Markov branching processes of Section 2.2 simplify the biology of growing cultures. Constant per-cell event rates imply exponentially distributed times between events, whereas age-dependent branching processes, which allow general distributions of cell-cycle duration, are often used for clonal data (Hyrien et al. 2010; Chen et al. 2011). A supercritical one-type age-dependent process also has a random establishment amplitude (Athreya and Ney 1972, Chapter IV, Section 11, Theorem 1). Extending the founder-aware likelihood to such processes requires a new derivation of the amplitude distribution and the matrix *B* in Equation (12c). The present derivation of *B* uses the infinitesimal covariance *D* of the Markov process. These processes omit density dependence and therefore describe cultures only while their growth is not limited by space or nutrients.

Even when the data follow the Markov branching model of Section 2.2 exactly, the founder-aware likelihood *M* involves several approximations. Matching two amplitude moments does not establish a Gamma density, and using eventual survivors to approximate finite-time survivors introduces an additional error. The Gaussian form of the conditional distribution of the population given the amplitude, and its centering at 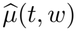, are also approximations. The numerical accuracy checks of Section 3.2 concern the computation of *M*, not these approximations. The Yule comparison shows early-time failure even with an exact amplitude distribution (Figure 1C). Finite-time corrections to the approximate conditional mean 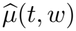 and covariance 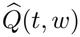 could improve the approximation at early times but are not developed here. The two-type experiments use one observation design and one set of true rates. For this setting, they measure the RMSE and coverage of *M* over repeated datasets, together with its mean log density relative to the other likelihoods. Because the exact observation density of the two-type model was not computed, they do not show how closely *M* approximates it.

Hypothesis (H4) also limits the rates to which the construction applies. In the two-type model, it fails when switching is much slower than net growth and the slower type grows at more than about half the net rate of the faster type (Section 2.3). The complementary component of *Z*(*t*) then has its own random limit, which is typically not Gaussian (Janson 2004, Theorem 3.1). A population founded by few cells therefore retains, in addition to its amplitude, a second random quantity from its early history, which records how early events shifted its composition away from the stable type proportions given by *r*. An extension to this regime requires the joint distribution of the amplitude and the second limit, together with a new residual approximation. In our experiments, switching rates comparable to the division rates keep (H4) satisfied throughout the fitting region (Section 3.1). In applications in which switching is slow compared with growth, (H4) should be verified over the fitting region before the founder-aware likelihood is used.

The construction of the founder-aware likelihood requires only the mean-growth matrix *A*, the infinitesimal covariance *D*(*z*), and the known founder state, together with hypotheses (H1)–(H4) of Section 2.3. Additional cell types, mixed founder states, intermediate founder counts, and observations that are linear functions of the counts therefore require no new derivation. Each new setting still requires verification of the hypotheses and an assessment of accuracy at finite times.

For an unknown number of founders, a problem also studied for quantitative polymerase chain reaction (Peccoud and Jacob 1996; Chigansky et al. 2018), a proposed extension is to average the likelihood over a seeding distribution, such as the Poisson distribution of cells plated at limiting dilution (Hu and Smyth 2009). Lineage-barcoding studies of single-founder clones (Mathis et al. 2017; Lan et al. 2017) require other observation models, such as sequencing read counts. Such studies may also show heritable differences in rates between clones (Mathis et al. 2017), whereas our model assumes that all cultures share the same rates. Because much of the variation in size among single-founder clones arises from their first few divisions, deaths, and switches alone (Equation (5)), the founder-aware likelihood could serve as a reference distribution under shared rates when testing for such differences.

Longitudinal observation of the same populations is a natural extension, for which Gunnarsson et al. (2023, Section 4.2.4) suggested developing a dedicated estimation method. Applied to longitudinal data, the founder-aware likelihood would share one latent amplitude among all observations of a population and would require the covariance of the residuals at different observation times conditional on that amplitude. A candidate approximation follows from the same decomposition of the residual that gives the matrix *B* (Online Resource 1, Section S1.2). This conditional dependence across times is what a fitted constant conditional variance cannot represent, so longitudinal data could separate *M* from *M*_const_ more sharply than the present design.

Two considerations may limit the advantage of the founder-aware likelihood for longitudinal data. With observations at several times, the growth between observations provides information about the rates, and the population size then determines the amplitude closely. The assumed shape of the amplitude distribution should therefore matter less than in endpoint designs. Once a population is large, conditioning on the previous observation also makes each subsequent step a many-founder problem. For single-founder populations observed longitudinally, Gaussian pseudo-likelihood conditional on previous counts performed substantially better than its unconditional form (Chen and Hyrien 2011, Table 2). Relevant comparators are these conditional likelihoods, composite likelihoods for longitudinal clonal data (Chen et al. 2011), and Gaussian approximations to demographic noise along observed trajectories, such as the linear noise approximation used in reaction-network inference (Fearnhead et al. 2014; Schnoerr et al. 2017). The accuracy of the founder-aware likelihood for longitudinal data, including the early-time error shown in Figure 1C, would need the same kind of assessment as in this study.

## 6 Conclusions

A population founded by few cells retains the effect of its earliest divisions, deaths, and switches however large it grows. This effect, summarized by the establishment amplitude, is shared by all descendants in a culture, so growth does not average it out; only additional independent founders do. A Gaussian likelihood built from the exact population mean and covariance can therefore misrepresent cultures founded by few cells even when each contains hundreds of cells, and rates estimated with it can appear more precise than the data support. The founder-aware likelihood treats the amplitude and the remaining fluctuations separately. It retains the amplitude distribution, which is far from Gaussian when founders are few. It approximates by a Gaussian distribution only the fluctuations that remain once the amplitude is specified. This approximation requires a large population rather than many founders. In our experiments, this separation improved rate estimation and the coverage of profile sets with one founder per culture and made no resolved difference with 50 founders. The founder-aware likelihood therefore suits clonal and other few-founder experiments (Hyrien et al. 2010; Mathis et al. 2017), whereas the cheaper population Gaussian likelihood *G* suffices when cultures can be seeded with many cells.

### Supplementary information

Online Resource 1 (ESM_1.pdf) derives the residual approximation and the amplitude moments, gives the survival probabilities of the two-type model and the error from substituting eventual survival for survival to the observation time, gives the exact reference distribution for the Yule process, and describes the numerical methods, accuracy checks, computational costs, and additional simulation results.

## Supporting information

Online Resource 1: Supplementary information

## Statements and Declarations

### Data and code availability

The study uses only synthetic data, generated by the simulation code from the experiment configurations and their random seeds. The code, the configurations, and the summary results from which all figures and tables are built are available at https://github.com/juhang62/founder-aware-likelihood. The repository also gives the commands that regenerate the simulated observations, fits, and profile sets.

### Author contributions

**Lifeng Han:** Conceptualization, Methodology, Software, Validation, Formal analysis, Investigation, Data curation, Visualization, Writing – original draft, Writing – review & editing, Supervision, Project administration. **Marley M. Assefa:** Conceptualization, Investigation, Writing – review & editing.

### Use of generative AI tools

OpenAI Codex was used to assist with organization, drafting, and code for figures generated from saved numerical summaries. Anthropic Claude was used to assist with revising the framing and drafting of the text.

## References

Asmussen, S., Hering, H.: Branching Processes. Progress in Probability and Statistics, vol. 3. Birkhäuser, Boston (1983). 10.1007/978-1-4615-8155-0

Athreya, K.B., Ney, P.E.: Branching Processes. Springer, Berlin (1972). 10.1007/978-3-642-65371-1

Athreya, K.B.: Some results on multitype continuous time Markov branching processes. Annals of Mathematical Statistics 39(2), 347–357 (1968) 10.1214/aoms/1177698395

Athreya, K.B.: Limit theorems for multitype continuous time Markov branching processes. II. The case of an arbitrary linear functional. Zeitschrift für Wahrschein-lichkeitstheorie und verwandte Gebiete 13, 204–214 (1969) 10.1007/BF00539201

Baker, J., Chigansky, P., Hamza, K., Klebaner, F.C.: Persistence of small noise and random initial conditions. Advances in Applied Probability 50(A), 67–81 (2018) 10.1017/apr.2018.71

Barbour, A.D., Chigansky, P., Klebaner, F.C.: On the emergence of random initial conditions in fluid limits. Journal of Applied Probability 53(4), 1193–1205 (2016) 10.1017/jpr.2016.74

Barbour, A.D., Hamza, K., Kaspi, H., Klebaner, F.C.: Escape from the boundary in Markov population processes. Advances in Applied Probability 47(4), 1190–1211 (2015) 10.1017/s0001867800049077

Cheek, D., Antal, T.: Mutation frequencies in a birth–death branching process. The Annals of Applied Probability 28(6), 3922–3947 (2018) 10.1214/18-AAP1413

Chen, R., Hyrien, O.: Quasi- and pseudo-maximum likelihood estimators for discretely observed continuous-time Markov branching processes. Journal of Statistical Planning and Inference 141(7), 2209–2227 (2011) 10.1016/j.jspi.2011.01.016

Chen, R., Hyrien, O., Noble, M., Mayer-Pröschel, M.: A composite likelihood approach to the analysis of longitudinal clonal data on multitype cellular systems under an age-dependent branching process. Biostatistics 12(1), 173–191 (2011) 10.1093/biostatistics/kxq050

Chigansky, P., Jagers, P., Klebaner, F.C.: What can be observed in real time PCR and when does it show? Journal of Mathematical Biology 76(3), 679–695 (2018) 10.1007/s00285-017-1154-1

Chigansky, P., Jagers, P., Klebaner, F.C.: Populations with interaction and environmental dependence: from few, (almost) independent, members into deterministic evolution of high densities. Stochastic Models 35(2), 108–118 (2019) 10.1080/15326349.2019.1575755

Clopper, C.J., Pearson, E.S.: The use of confidence or fiducial limits illustrated in the case of the binomial. Biometrika 26(4), 404–413 (1934) 10.1093/biomet/26.4.404

Davison, A.C., Hautphenne, S., Kraus, A.: Parameter estimation for discretely observed linear birth-and-death processes. Biometrics 77(1), 186–196 (2021) 10.1111/biom.13282

Durrett, R.: Branching Process Models of Cancer. Springer, Cham (2015). 10.1007/978-3-319-16065-8

Fearnhead, P., Giagos, V., Sherlock, C.: Inference for reaction networks using the linear noise approximation. Biometrics 70, 457–466 (2014) 10.1111/biom.12152

Franken, N.A.P., Rodermond, H.M., Stap, J., Haveman, J., Bree, C.: Clonogenic assay of cells in vitro. Nature Protocols 1(5), 2315–2319 (2006) 10.1038/nprot.2006.339

Georgii, H.-O., Baake, E.: Supercritical multitype branching processes: the ancestral 26 types of typical individuals. Advances in Applied Probability 35(4), 1090–1110 (2003) 10.1239/aap/1067436336

Gunnarsson, E.B., De, S., Leder, K., Foo, J.: Understanding the role of phenotypic switching in cancer drug resistance. Journal of Theoretical Biology 490, 110162 (2020) 10.1016/j.jtbi.2020.110162

Gupta, P.B., Fillmore, C.M., Jiang, G., Shapira, S.D., Tao, K., Kuperwasser, C., Lander, E.S.: Stochastic state transitions give rise to phenotypic equilibrium in populations of cancer cells. Cell 146(4), 633–644 (2011) 10.1016/j.cell.2011.07.026

Gunnarsson, E.B., Foo, J., Leder, K.: Statistical inference of the rates of cell proliferation and phenotypic switching in cancer. Journal of Theoretical Biology 568, 111497 (2023) 10.1016/j.jtbi.2023.111497

Gillespie, D.T.: Exact stochastic simulation of coupled chemical reactions. The Journal of Physical Chemistry 81(25), 2340–2361 (1977) 10.1021/j100540a008

Gourieroux, C., Monfort, A., Trognon, A.: Pseudo maximum likelihood methods: theory. Econometrica 52(3), 681–700 (1984) 10.2307/1913471

Hyrien, O., Chen, R., Mayer-Pröschel, M., Noble, M.: Saddlepoint approximations to the moments of multitype age-dependent branching processes, with applications. Biometrics 66(2), 567–577 (2010) 10.1111/j.1541-0420.2009.01281.x

Hu, Y., Smyth, G.K.: ELDA: extreme limiting dilution analysis for comparing depleted and enriched populations in stem cell and other assays. Journal of Immunological Methods 347(1–2), 70–78 (2009) 10.1016/j.jim.2009.06.008

Ho, L.S.T., Xu, J., Crawford, F.W., Minin, V.N., Suchard, M.A.: Birth/birth-death processes and their computable transition probabilities with biological applications. Journal of Mathematical Biology 76(4), 911–944 (2018) 10.1007/s00285-017-1160-3

Janson, S.: Functional limit theorems for multitype branching processes and generalized Pólya urns. Stochastic Processes and their Applications 110, 177–245 (2004) 10.1016/j.spa.2003.12.002

Kimmel, M., Axelrod, D.E.: Branching Processes in Biology, 2nd edn. Interdisciplinary Applied Mathematics. Springer, New York (2015). 10.1007/978-1-4939-1559-0

Kendall, D.G.: Branching processes since 1873. Journal of the London Mathematical Society s1-41(1), 385–406 (1966) 10.1112/jlms/s1-41.1.385

Kent, J.T.: Robust properties of likelihood ratio tests. Biometrika 69(1), 19–27 (1982) 10.1093/biomet/69.1.19

Kolesko, K., Kołodziejska, A., Meiners, M.: Asymptotic fluctuations in supercritical multi-type Crump–Mode–Jagers processes. arXiv preprint arXiv:2607.22414v1 (2026). https://arxiv.org/abs/2607.22414v1

Kesten, H., Stigum, B.P.: Additional limit theorems for indecomposable multidimensional Galton–Watson processes. Annals of Mathematical Statistics 37(6), 1463–1481 (1966) 10.1214/aoms/1177699139

Kesten, H., Stigum, B.P.: A limit theorem for multidimensional Galton–Watson processes. The Annals of Mathematical Statistics 37(5), 1211–1223 (1966) 10.1214/aoms/1177699266

Lan, X., Jörg, D.J., Cavalli, F.M.G., Richards, L.M., Nguyen, L.V., Vanner, R.J., Guilhamon, P., Lee, L., Kushida, M.M., Pellacani, D., Park, N.I., Coutinho, F.J., et al.: Fate mapping of human glioblastoma reveals an invariant stem cell hierarchy. Nature 549(7671), 227–232 (2017) 10.1038/nature23666

Luo, X.G., Kuipers, J., Beerenwinkel, N.: Numerical approximations of population size distributions for multi-type branching processes. arXiv preprint arXiv:2609.07526v1 (2026). https://arxiv.org/abs/2609.07526v1

Morris, D.J., Kennedy, L., Black, A.J.: Random time-shift approximation enables hierarchical Bayesian inference of mechanistic within-host viral dynamics models on large datasets. PLOS Computational Biology 21, 1013775 (2025) 10.1371/journal.pcbi.1013775

Morris, D., Maclean, J., Black, A.J.: Computation of random time-shift distributions for stochastic population models. Journal of Mathematical Biology 89, 33 (2024) 10.1007/s00285-024-02132-6

Mathis, R.A., Sokol, E.S., Gupta, P.B.: Cancer cells exhibit clonal diversity in phenotypic plasticity. Open Biology 7(2), 160283 (2017) 10.1098/rsob.160283

Piessens, R., Doncker-Kapenga, E., Überhuber, C.W., Kahaner, D.K.: QUADPACK: A Subroutine Package for Automatic Integration. Springer Series in Computational Mathematics, vol. 1. Springer, Berlin, Heidelberg (1983). 10.1007/978-3-642-61786-7

Peccoud, J., Jacob, C.: Theoretical uncertainty of measurements using quantitative polymerase chain reaction. Biophysical Journal 71(1), 101–108 (1996) 10.1016/S0006-3495(96)79205-6

Russo, M., Pompei, S., Sogari, A., Corigliano, M., Crisafulli, G., Puliafito, A., Lamba, S., Erriquez, J., Bertotti, A., Gherardi, M., Di Nicolantonio, F., Bardelli, A., et al.: A modified fluctuation-test framework characterizes the population dynamics and mutation rate of colorectal cancer persister cells. Nature Genetics 54(7), 976–984 (2022) 10.1038/s41588-022-01105-z

Ren, Y.-X., Song, R., Zhang, R.: Central limit theorems for supercritical branching Markov processes. Journal of Functional Analysis 266(3), 1716–1756 (2014) 10.1016/j.jfa.2013.10.015

Ren, Y.-X., Song, R., Zhang, R.: Central limit theorems for supercritical branching nonsymmetric Markov processes. Annals of Probability 45(1), 564–623 (2017) 10.1214/14-AOP987

Shaffer, S.M., Dunagin, M.C., Torborg, S.R., Torre, E.A., Emert, B., Krepler, C., Beqiri, M., Sproesser, K., Brafford, P.A., Xiao, M., Eggan, E., Anastopoulos, I.N., et al.: Rare cell variability and drug-induced reprogramming as a mode of cancer drug resistance. Nature 546(7658), 431–435 (2017) 10.1038/nature22794

Sharma, S.V., Lee, D.Y., Li, B., Quinlan, M.P., Takahashi, F., Maheswaran, S., McDermott, U., Azizian, N., Zou, L., Fischbach, M.A., Wong, K.-K., Brandstetter, K., et al.: A chromatin-mediated reversible drug-tolerant state in cancer cell subpopulations. Cell 141(1), 69–80 (2010) 10.1016/j.cell.2010.02.027

Schnoerr, D., Sanguinetti, G., Grima, R.: Approximation and inference methods for stochastic biochemical kinetics—a tutorial review. Journal of Physics A: Mathematical and Theoretical 50(9), 093001 (2017) 10.1088/1751-8121/aa54d9

Virtanen, P., Gommers, R., Oliphant, T.E., Haberland, M., Reddy, T., Cournapeau, D., Burovski, E., Peterson, P., Weckesser, W., Bright, J., Walt, S.J., Brett, M., et al.: SciPy 1.0: fundamental algorithms for scientific computing in Python. Nature Methods 17(3), 261–272 (2020) 10.1038/s41592-019-0686-2

Wu, C., Gunnarsson, E.B., Foo, J., Leder, K.: A statistical framework for detecting therapy-induced resistance from drug screens. npj Systems Biology and Applications 11, 88 (2025) 10.1038/s41540-025-00560-8

Wu, C., Gunnarsson, E.B., Myklebust, E.M., Köhn-Luque, A., Tadele, D.S., Enserink, J.M., Frigessi, A., Foo, J., Leder, K.: Using birth-death processes to infer tumor subpopulation structure from live-cell imaging drug screening data. PLOS Computational Biology 20(3), 1011888 (2024) 10.1371/journal.pcbi.1011888

Xu, J., Guttorp, P., Kato-Maeda, M., Minin, V.N.: Likelihood-based inference for discretely observed birth–death-shift processes, with applications to evolution of mobile genetic elements. Biometrics 71(4), 1009–1021 (2015) 10.1111/biom.12352

Yakovlev, A.Y., Yanev, N.M.: Relative frequencies in multitype branching processes. The Annals of Applied Probability 19(1), 1–14 (2009) 10.1214/08-AAP539

