## Supplementary material for "Large populations from few founders: founder-aware likelihoods for inferring cell proliferation rates": Online Resource 1: Supplementary information

Supplementary information for an article in *Bulletin of Mathematical Biology*

Lifeng Han* Marley M. Assefa

Department of Mathematics, Tulane University, 6823 St. Charles Avenue, New Orleans, 70118, USA

This supplement provides mathematical derivations and numerical details for the likelihoods compared in the main text. It uses the notation of the main text and restates each definition, or cites the main-text equation that gives it, where the quantity first appears. The labels $G$, $EG$, $M$, $M_0$, and $M_{\text{const}}$ denote the five likelihoods of Table 1 of the main text: the population Gaussian likelihood, its adaptation with a separate empty-culture component, the founder-aware likelihood, and the two variants of the founder-aware likelihood that set the approximate conditional covariance $\widehat{Q}(t, w)$ to zero or replace it by a fitted constant conditional variance.

Section S1 develops the residual approximation and the amplitude moments for $q$ cell types, with a specialization to the two-type model. It begins with the martingale tools used throughout (Section S1.1) and states the residual limit theorem (Section S1.2). The second moments derived in Section S1.2 identify the matrix $B$ in the approximate conditional covariance $\widehat{Q}(t, w) = w \exp(Lt)B$. Two ways of computing $B$ and the derivation of the amplitude moments follow in Sections S1.3 and S1.4, respectively. For the two-type model, Section S2 verifies the hypotheses over the fitting region and calculates the survival probabilities. It also bounds the error from approximating the distribution of cultures alive at the observation time by that of cultures that survive indefinitely.

Section S3 gives the exact distribution of the one-type Yule process used as the reference in Figure 1 of the main text. Section S4 describes the numerical methods and reports the accuracy checks and computational times. Section S5 gives additional simulation results, including uncertainty in paired comparisons of the likelihoods. These numerical methods and additional results concern the two-type model. Numerical entries come from the experiments described in the main text; no additional simulations or parameter fits were performed for this supplement.

### S1 Residual approximation and amplitude moments

#### S1.1 Model, hypotheses, and martingale tools

Let $Z(t) = (Z_1(t), \ldots, Z_q(t))^\top$ be the column vector of counts of $q$ cell types at time $t \geq 0$, started from a known nonzero initial state $z_0$. As in Section 2.2 of the main text, each type-$j$ cell evolves independently of all others and can undergo finitely many kinds of events. An event of kind $k$ occurs at constant rate $\alpha_{jk}$ per cell and changes the vector of counts by a bounded integer vector $\nu_{jk}$. With $e_j$ the $j$th coordinate vector, the mean-growth matrix $A$ and the infinitesimal covariance $D(z)$ at state $z = (z_1, \ldots, z_q)^\top$ are given by Equation (6) of the main text,

$$Ae_j = \sum_k \alpha_{jk}\nu_{jk}, \qquad D(z) = \sum_{j=1}^{q} z_j \sum_k \alpha_{jk}\nu_{jk}\nu_{jk}^\top.$$

The map $z \mapsto D(z)$ is linear, and $D(z)$ is positive semidefinite whenever $z$ has nonnegative entries. We use hypotheses (H1)–(H4) of Section 2.3 of the main text. Hypothesis (H1) is the model just described, with finite event rates. Hypothesis (H2) requires that $A$ be irreducible, so that every type can eventually produce every other type. Hypothesis (H3) requires that the dominant eigenvalue $L$ of $A$ be positive. Hypothesis (H4) requires that every other eigenvalue $\lambda$ of $A$ satisfy $\operatorname{Re}\lambda < L/2$. Under (H2), the right and left eigenvectors $r$ and $\ell$ of $A$ for $L$ have positive entries, and we normalize them by $r_q = 1$ and $\ell^\top r = 1$.

Let $P = r\ell^\top$ and $S = I_q - P$, with $I_q$ the $q \times q$ identity matrix. Since $Ar = Lr$, $\ell^\top A = L\ell^\top$, and $\ell^\top r = 1$, we have $P^2 = P$ and $AP = PA = LP$, so $S^2 = S$ and $AS = SA$. Thus $P$ and $S$ are the spectral projections of $A$ for its dominant eigenvalue $L$ and for its remaining eigenvalues. Consequently, $\exp(At)P = \exp(Lt)P$, and the entries of $\exp(At)S$ are linear combinations of the functions $t^j \exp(\lambda t)$, where $\lambda$ is an eigenvalue of $A$ other than $L$ and $j$ is a nonnegative integer smaller than the size of a Jordan block for $\lambda$. In the two-type model, the other eigenvalue $\lambda_2$ is real and distinct from $L$ because both switching rates are positive. Consequently, $\exp(At)S = \exp(\lambda_2 t)S$. The entries of $A$ are given in Section S2.1 and Equation (7) of the main text.

Write $K = A - (L/2)I_q$, and let $\|\cdot\|$ denote the Euclidean norm of vectors and the induced norm of matrices. The matrix $\exp(Kx)S$ decays exponentially in $x$, as follows. For $q > 1$, choose and fix

$$0 < \rho < \frac{L}{2} - \max_{\lambda \neq L} \operatorname{Re}\lambda,$$

where the maximum is over the eigenvalues of $A$ other than $L$. This interval is nonempty by (H4). Since $\exp(Kx)S = \exp(-Lx/2)\exp(Ax)S$, the entries of $\exp(Kx)S$ are linear combinations of the functions $x^j \exp((\lambda - L/2)x)$, with $\lambda$ and $j$ as in the previous paragraph. Since $L/2 - \operatorname{Re}\lambda - \rho > 0$, each factor $x^j \exp(-(L/2 - \operatorname{Re}\lambda - \rho)x)$ is bounded for $x \geq 0$. Consequently, there is a constant $\gamma_0 < \infty$, depending on the model and the chosen $\rho$, such that

$$\|\exp(Kx)S\| \leq \gamma_0 \exp(-\rho x), \qquad x \geq 0. \tag{S1}$$

When $q = 1$, $S = 0$, so the bound holds for any $\rho > 0$.

Let $(\mathcal{F}_t)_{t\geq 0}$ be the completed, right-continuous filtration generated by $Z$, where $\mathcal{F}_t$ represents the information available from the population history up to time $t$. The compensated population martingale is

$$\mathcal{M}(t) = Z(t) - z_0 - \int_0^t AZ(u)\,du. \tag{S2}$$

To calculate the second moments of the martingale, we use the predictable quadratic covariation matrix, also called the bracket. We write $\langle\mathcal{M}\rangle$ for the bracket process and $\langle\mathcal{M}\rangle_t$ for its value at time $t$; the same convention applies to its scalar entries. For this model it is

$$\langle\mathcal{M}\rangle_t = \int_0^t D(Z(u))\,du. \tag{S3}$$

Each possible population jump contributes its outer product multiplied by its current event rate. Summing these contributions gives $D(Z(u))$. The bracket therefore accumulates the infinitesimal covariance along the population path. It is continuous and depends only on the population history, which ensures predictability here.

The bracket is characterized by the requirement that $\mathcal{M}_i(t)\mathcal{M}_j(t) - \langle\mathcal{M}_i, \mathcal{M}_j\rangle_t$ be a martingale for each pair of components $i, j$ (Protter 2005, Chapter III, Section 5, pp. 124–125). Taking expectations gives $\mathbb{E}\langle\mathcal{M}\rangle_t = \operatorname{Cov}(\mathcal{M}(t))$, although the bracket itself is generally random.

Define $W_t = \exp(-Lt)\ell^\top Z(t)$ and $a_0 = \ell^\top z_0$, which is positive because $\ell$ has positive entries and $z_0$ is a nonzero vector of counts. Because $Z$ has paths of finite variation and $\exp(-Lt)$ is deterministic and continuously differentiable, the product rule, Equation (S2), and $\ell^\top A = L\ell^\top$ give

$$W_t = a_0 + \int_0^t \exp(-Lu)\ell^\top \, d\mathcal{M}(u). \tag{S4}$$

Thus $W_t$, which we call the amplitude martingale, is a martingale. Under (H1)–(H3), it converges almost surely and in mean square as $t \to \infty$ (Janson 2004, Lemma 9.5). Its limit $W = \lim_{t\to\infty} W_t$ is the establishment amplitude of Equation (10) of the main text. Its mean is $\mathbb{E}W = a_0$ (Section S1.4), and $W > 0$ exactly on eventual survival, up to null events. The residual at time $t$ is $U(t) = Z(t) - W\exp(Lt)r$, as defined in Equation (11) of the main text. This pathwise definition does not condition on $W$ or imply a zero conditional mean.

Let $\Psi$ be a deterministic continuous matrix-valued function and $I \subset [0,\infty)$ a bounded interval. Because $\mathcal{M}$ is a square-integrable martingale and $\Psi$ is deterministic, the stochastic integral $\int_I \Psi \, d\mathcal{M}$ has mean zero. Its second moment is given by the isometry

$$\mathbb{E}\Big[\Big(\int_I \Psi \, d\mathcal{M}\Big)\Big(\int_I \Psi \, d\mathcal{M}\Big)^\top\Big] = \int_I \Psi(u)\, D(m(u))\, \Psi(u)^\top du, \tag{S5}$$

where $m(u) = \mathbb{E}Z(u) = \exp(Au)z_0$ is the population mean, as in the main text. To obtain Equation (S5), we use the bracket in Equation (S3) and the quadratic covariation formula for stochastic integrals. We then take expectations using the defining martingale property of the predictable bracket (Protter 2005, Chapter II, Sections 5–6, and Chapter III, Section 5) and the identity $\mathbb{E}D(Z(u)) = D(m(u))$, which follows from the linearity of $D$.

#### S1.2 The joint limit and second moments of the residual

Theorem 1 of the main text states that, under (H1)–(H4) and conditional on eventual survival $W > 0$, the pair $(W, \exp(-Lt/2)U(t))$ converges in distribution to $(W, \sqrt{W}\,\xi)$ as $t \to \infty$, where $\xi$ is a $q$-dimensional Gaussian random vector with mean zero and covariance $B$, independent of $W$. The same limit holds with $U(t)$ replaced by $Z(t) - \widehat{\mu}(t, W)$, using the centering $\widehat{\mu}(t, W)$ defined below. We cite this theorem from published work and describe the scope of each citation below:

Asmussen and Hering (1983, Chapter VIII, Corollary 3.2 and Theorem 12.1) stated the limit of each linear combination of the counts conditional on survival, for general multitype processes in discrete and continuous time, without proof. Ren et al. (2014, Theorem 1.10) and Ren et al. (2017, Theorem 1.16 and Remark 1.21) proved the joint limit with $W$, with the Gaussian factor independent of $W$, for any initial population when cells change type only by switching. Kolesko et al. (2026, Theorem 2.8(i) and Remark 2.11) proved it for general multitype branching processes started from one individual; the extension to several founders follows because their lineages are independent. The decomposition into dominant and complementary directions and the limiting covariance in the complementary directions are established in branching fluctuation theory (Janson 2004). The amplitude martingale and its convergence are classical (Athreya and Ney 1972).

This subsection computes the second moments of the scaled residual under the original branching law, without conditioning on amplitude or survival. They identify the matrix $B$ in the joint limiting law, under which $\sqrt{W}\,\xi$ has covariance $wB$ conditional on $W = w$. Assume (H1)–(H4) and a fixed nonzero initial state $z_0$, and write $X_t = \exp(-Lt/2)U(t)$ for the scaled residual. We derive an exact decomposition of the residual, define $B$, and show that $\mathbb{E}\, X_t X_t^\top$ converges to $a_0 B$.

Variation of constants in Equation (S2), using $AS = SA$, gives the complementary component

$$SZ(t) = \exp(At)Sz_0 + \int_0^t \exp(A(t-u))S\,d\mathcal{M}(u).$$

For the dominant component, $P = r\ell^\top$ and $\ell^\top A = L\ell^\top$ give

$$PZ(t) = \exp(Lt)W_t r, \qquad W - W_t = \int_t^\infty \exp(-Lu)\ell^\top\,d\mathcal{M}(u).$$

For $T > t$, Equation (S4) gives $W_T - W_t = \int_t^T \exp(-Lu)\ell^\top\,d\mathcal{M}(u)$, and the second identity follows by taking the mean-square limit as $T \to \infty$. This limit defines the stochastic integral over $(t, \infty)$. Since $Z(t) = SZ(t) + PZ(t)$, adding the complementary component and $\exp(Lt)(W_t - W)r$ yields the exact decomposition

$$\begin{aligned} U(t) = \; & \exp(At)Sz_0 + \int_0^t \exp(A(t-u))S\,d\mathcal{M}(u) \\ & - \exp(Lt)\int_t^\infty \exp(-Lu)P\,d\mathcal{M}(u). \end{aligned} \tag{S6}$$

We call the integral over $[0, t]$ in Equation (S6) the past integral and the integral over $(t, \infty)$ the future integral. The future integral appears because the centering uses the limit $W$ rather than the current value $W_t$. The future integral in Equation (S6) is the mean-square limit of integrals over $(t, T]$ as $T \to \infty$, so Equation (S5) also holds on $(t, \infty)$. This integral has conditional mean zero given $\mathcal{F}_t$, whereas the deterministic term and the past integral are $\mathcal{F}_t$-measurable, so the three terms of Equation (S6) are uncorrelated.

Define

$$C_S = \int_0^\infty \exp(Ku)SD(r)S^\top \exp(K^\top u)\,du. \tag{S7}$$

Under (H4), the integral is finite because $\|\exp(Ku)S\|$ decays exponentially in $u$ (Equation (S1)). After multiplication by $\exp(-Lt/2)$, the second moments of the past and future integrals in Equation (S6) converge to $a_0\,C_S$ and $a_0\,PD(r)P^\top/L$, as shown below. The matrix $B$ of Equation (12c) of the main text is the sum of $C_S$ and $PD(r)P^\top/L$. In the founder-aware likelihood, we use this matrix to approximate the covariance of $U(t)$ conditional on $W = w$ by $\widehat{Q}(t, w)$ (Equation (12b) of the main text):

$$B = \frac{PD(r)P^\top}{L} + C_S, \qquad \widehat{Q}(t, w) = w\exp(Lt)B. \tag{S8}$$

Both terms on the right side of the expression for $B$ are positive semidefinite: $D(r)$ is positive semidefinite because $r$ has positive entries, so $PD(r)P^\top/L$ is positive semidefinite, and $C_S$ is the integral of the positive semidefinite matrices $\exp(Ku)SD(r)\big(\exp(Ku)S\big)^\top$.

To compute the second moments, we multiply Equation (S6) by $\exp(-Lt/2)$ and apply Equation (S5) to the two integrals. We then substitute $u = t - v$ in the integral over past times and $u = t + v$ in the integral over future times. Using $\exp(-Lv/2)\exp(Av) = \exp(Kv)$ and the linearity of $D$ gives the exact identity

$$\begin{aligned} \mathbb{E}\, X_t X_t^\top = \; & \exp(Kt)Sz_0 z_0^\top S^\top \exp(K^\top t) \\ & + \int_0^t \exp(Kv)S\,D\big(\exp(-L(t-v))m(t-v)\big)S^\top \exp(K^\top v)\,dv \\ & + \int_0^\infty \exp(-Lv)P\,D\big(\exp(-L(t+v))m(t+v)\big)P^\top dv. \end{aligned} \tag{S9}$$

This identity gives the second-moment matrix of the scaled residual at each finite time. Its covariance is obtained by subtracting the first term, the outer product of its mean $\mathbb{E}X_t = \exp(Kt)Sz_0$.

The scaled mean satisfies

$$\exp(-Ls)m(s) = a_0 r + \exp(-Ls/2)\exp(Ks)Sz_0,$$

because $Pz_0 = a_0 r$ and $\exp(-Ls/2)\exp(As)S = \exp(Ks)S$; it is therefore bounded for $s \geq 0$ and converges to $a_0 r$ as $s \to \infty$.

We now let $t \to \infty$ in each term of Equation (S9). By Equation (S1), the first term has norm at most $\gamma_0^2 \exp(-2\rho t)\|z_0\|^2$ and tends to zero. In the two integrals, take the integrand of the past integral to be zero for $v > t$, so that both are integrals over $v \geq 0$. For each fixed $v$, the scaled means $\exp(-L(t-v))m(t-v)$ and $\exp(-L(t+v))m(t+v)$ in the arguments of $D$ converge to $a_0 r$. By the linearity of $D$, the two integrands therefore converge to $a_0 \exp(Kv)SD(r)S^\top \exp(K^\top v)$ and $a_0 \exp(-Lv)PD(r)P^\top$, respectively. Because the scaled mean is bounded and $D$ is linear, Equation (S1) also bounds the norms of the two integrands, for every $t$, by constant multiples of $\exp(-2\rho v)$ and $\exp(-Lv)$, which are integrable over $v \geq 0$. Lebesgue's dominated convergence theorem therefore allows the limit $t \to \infty$ to be taken inside each integral. With Equation (S7) and $\int_0^\infty \exp(-Lv)\,dv = 1/L$, this gives

$$\lim_{t\to\infty} \mathbb{E}\, X_t X_t^\top = a_0\, C_S + a_0 \frac{PD(r)P^\top}{L} = a_0\, B. \tag{S10}$$

On extinction, the scaled residual is eventually zero, because $Z(t)$ is eventually zero and $W = 0$. Conditional on survival, it converges in distribution to $\sqrt{W}\,\xi$ by Theorem 1. Combining these two cases gives convergence in distribution to $\sqrt{W}\,\xi$ under the original branching law, with $W = 0$ on extinction. Because $\xi$ has covariance $B$ and is independent of $W$, this limit has second moment $\mathbb{E}[W]B = a_0 B$, in agreement with Equation (S10). Equating the two limits requires, in addition, uniform integrability of $\|X_t\|^2$, which follows from a uniform bound on fourth moments for processes with bounded jumps. We do not reproduce that argument here.

At a specified amplitude $W = w$, the founder-aware likelihood centers its Gaussian component at $\widehat{\mu}(t, w) = w\, m(t)/a_0$ rather than at $w \exp(Lt) r$ (Equation (12a) of the main text), where $m(t) = \mathbb{E}Z(t)$ is the population mean. The corresponding approximation to the conditional distribution of $U(t)$ has mean $\Delta(t, w) = \widehat{\mu}(t, w) - w \exp(Lt) r$ and covariance $\widehat{Q}(t, w)$, as described after Equation (13) of the main text. Since $Pz_0 = a_0 r$,

$$\exp(-Lt/2)\Delta(t, w) = \frac{w}{a_0} \exp(Kt)Sz_0 \longrightarrow 0 \tag{S11}$$

for each fixed $w > 0$, in ordinary convergence of deterministic vectors. Equation (S11) shows that the adjustment $\Delta(t, w)$ is negligible on the fluctuation scale $\exp(Lt/2)$. The second statement of Theorem 1 follows from Equation (S11) with $w = W$:

$$\exp(-Lt/2)\Delta(t, W) = \frac{W}{a_0} \exp(Kt)Sz_0$$

tends to zero almost surely, so by Slutsky's lemma the same limit holds with $U(t)$ replaced by $U(t) - \Delta(t, W) = Z(t) - \widehat{\mu}(t, W)$.

The second moments above are computed under the original branching law, not conditional on a specified amplitude $W = w$; they identify $B$ in the joint limit but are not the conditional covariance of $U(t)$ given $W = w$. Because the founder-aware likelihood integrates over $W$, it uses only the joint limit of Theorem 1 stated in the main text. The finite-time accuracy of using this limit as an approximation is assessed numerically, as explained in Section 2.3 of the main text.

#### S1.3 Computing $B$

Equation (S8) defines $B$ through the integral $C_S$ of Equation (S7). We evaluate that integral in closed form when $A$ is diagonalizable and give a linear equation for $B$ that holds for any $A$ satisfying (H1)–(H4).

Suppose that $A$ is diagonalizable, with eigenvalues $\lambda_1 = L, \lambda_2, \ldots, \lambda_q$ and right and left eigenvectors $r_k$ and $\ell_k$. We normalize them by $\ell_j^\top r_k = \delta_{jk}$, where $\delta_{jk}$ is the Kronecker delta, and take $r_1 = r$ and $\ell_1 = \ell$. With $P_k = r_k \ell_k^\top$, we have $S = \sum_{k\geq 2} P_k$ and $K^n P_k = (\lambda_k - L/2)^n P_k$ for every $n \geq 0$. Substituting these identities into the power series $\exp(Ku) = \sum_{n\geq 0}(Ku)^n/n!$ that defines the matrix exponential gives $\exp(Ku)S = \sum_{k\geq 2} \exp((\lambda_k - L/2)u)P_k$. Substituting this into Equation (S7) and integrating each term gives

$$C_S = \sum_{j=2}^{q}\sum_{k=2}^{q} \frac{P_j D(r) P_k^\top}{L - \lambda_j - \lambda_k}, \tag{S12}$$

where each integral is finite because $\mathrm{Re}(\lambda_j + \lambda_k) < L$. Complex eigenvalues occur in conjugate pairs with conjugate projections, so the sum is real. In the two-type model, $A$ is diagonalizable because $\lambda_2$ is real and distinct from $L$, and only $j = k = 2$ contributes:

$$C_S = \frac{SD(r)S^\top}{L - 2\lambda_2}, \qquad B = \frac{PD(r)P^\top}{L} + \frac{SD(r)S^\top}{L - 2\lambda_2}, \tag{S13}$$

provided $\lambda_2 < L/2$. Our computations use this two-type form.

For any $q$, $B$ can also be computed by solving one linear system, without requiring $A$ to be diagonalizable. Define $\sigma^2 = \ell^\top D(r)\ell/L$. This scalar is positive. Indeed, since $\ell^\top A = L\ell^\top$, the definition $Ae_j = \sum_k \alpha_{jk}\nu_{jk}$ gives $\sum_k \alpha_{jk}\ell^\top \nu_{jk} = L\ell_j > 0$ for each type $j$. Thus, some event with positive rate has $\ell^\top \nu_{jk} \neq 0$, and hence $\ell^\top D(r)\ell = \sum_j r_j \sum_k \alpha_{jk}(\ell^\top \nu_{jk})^2 > 0$. Define

$$F = K - \frac{D(r)\ell\ell^\top}{\sigma^2}. \tag{S14}$$

Then $B$ is the unique solution of the Lyapunov equation

$$FB + BF^\top + D(r) = 0. \tag{S15}$$

We first derive an equation for $B$ that involves $K$ in place of $F$. By the product rule, the integrand of Equation (S7) satisfies

$$\begin{aligned}\frac{d}{du}\Big(\exp(Ku)SD(r)S^\top \exp(K^\top u)\Big) &= K\exp(Ku)SD(r)S^\top \exp(K^\top u) \\ &\quad + \exp(Ku)SD(r)S^\top \exp(K^\top u)K^\top.\end{aligned}$$

Integrate both sides from 0 to $\infty$. The integrand equals $SD(r)S^\top$ at $u = 0$ and tends to zero as $u \to \infty$ by Equation (S1), so the left side gives $-SD(r)S^\top$, and the right side gives $KC_S + C_S K^\top$. Write $C_P = PD(r)P^\top/L$ for the other term of $B$. Since $KP = (L/2)P$, we have $KC_P + C_P K^\top = LC_P = PD(r)P^\top$. Adding the two identities gives

$$KB + BK^\top = PD(r)P^\top - SD(r)S^\top. \tag{S16}$$

The added term in $F$ converts the right side of Equation (S16) into $-D(r)$. To evaluate that term, note that $C_P = \sigma^2 rr^\top$ and that $C_S\ell = 0$, because $S^\top \exp(K^\top u)\ell = \exp(Lu/2)S^\top\ell = 0$ for every $u$. Hence $B\ell = \sigma^2 r$ and, since $B$ is symmetric, $\ell^\top B = \sigma^2 r^\top$. It follows that $D(r)\ell\ell^\top B/\sigma^2 = D(r)\ell r^\top = D(r)P^\top$ and $B\ell\ell^\top D(r)/\sigma^2 = r\ell^\top D(r) = PD(r)$, so Equation (S16) gives

$$FB + BF^\top = PD(r)P^\top - SD(r)S^\top - D(r)P^\top - PD(r).$$

Since $P + S = I_q$, we have $D(r)P^\top = PD(r)P^\top + SD(r)P^\top$ and $PD(r) = PD(r)P^\top + PD(r)S^\top$, so the right side equals $-(P + S)D(r)(P + S)^\top = -D(r)$, which proves Equation (S15). The identity $B\ell = \sigma^2 r$ also shows that $\sigma^2 = \ell^\top B\ell$ is the variance of $\ell^\top\xi$, the component along $\ell$ of the limiting Gaussian vector $\xi$ of Section S1.2.

For uniqueness, note that $\ell^\top F = (L/2 - L)\ell^\top = -(L/2)\ell^\top$ and that $Fx = Kx$ whenever $\ell^\top x = 0$. Thus $F$ maps the range of $S$, which is $\{x : \ell^\top x = 0\}$, into itself and agrees with $K$ there. The eigenvalues of $F$ are therefore $-L/2$ and $\lambda_k - L/2$ for $k \geq 2$, counted with algebraic multiplicity, and all have negative real part by (H3) and (H4). The eigenvalues of the linear map $X \mapsto FX + XF^\top$ are sums of two eigenvalues of $F$, so none is zero. The map is therefore invertible, and Equation (S15) has exactly one solution. Equation (S16) also determines $B$ whenever the map $X \mapsto KX + XK^\top$ is invertible. The eigenvalues of that map include $(L - L/2) + (\lambda_k - L/2) = \lambda_k$ for each $k \geq 2$. The map is therefore singular at parameter values where $A$ has a zero eigenvalue and ill-conditioned near those values. Equation (S15), however, has a unique solution wherever (H1)–(H4) hold.

##### S1.4 Amplitude moments

Exact amplitude moments follow from the quadratic variation of the amplitude martingale. For the coordinate vectors $e_i$, let $c_i = \ell^\top D(e_i)\ell$ and $c = (c_1, \ldots, c_q)^\top$, and define $\eta = (\eta_1, \ldots, \eta_q)^\top$ by

$$(2LI_q - A^\top)\eta = c. \tag{S17}$$

Then $\eta_i$ is the amplitude variance from one type-$i$ founder, and $v_0 = \mathrm{Var}(W) = z_0^\top\eta$. Indeed, Equation (S4), the isometry in Equation (S5), and the mean-square convergence of $W_t$ to $W$ give $\mathbb{E}W = a_0$ and

$$v_0 = \lim_{t\to\infty} \mathbb{E}(W_t - a_0)^2 = \int_0^\infty \exp(-2Lu)\ell^\top D(\exp(Au)z_0)\ell\, du = z_0^\top(2LI_q - A^\top)^{-1}c. \tag{S18}$$

Together with $a_0 = \ell^\top z_0$ and the eventual survival probability $\pi_\infty = \Pr(W > 0)$, these moments give the moments of $W$ conditional on $W > 0$ in Equation (15) of the main text, which determine the Gamma approximation to the distribution of the positive amplitude.

#### S2 Two-type survival probabilities and the eventual-survival substitution

##### S2.1 The two-type model and hypotheses over the fitting region

This section concerns the two-type branching model. Each type-$i$ cell divides into two type-$i$ cells at rate $b_i$, dies at rate $d_i$, and switches to type $j \neq i$ at rate $s_{ij}$. The initial state is $z_0 = (n_1, n_2)^\top$, with $n_i$ founders of type $i$ and total founder count $N_0 = n_1 + n_2$. By Equation (7) of the main text, the diagonal entries of $A$ are $A_{11} = b_1 - d_1 - s_{12}$ and $A_{22} = b_2 - d_2 - s_{21}$, and its off-diagonal entries are $A_{12} = s_{21}$ and $A_{21} = s_{12}$.

The birth–death experiment uses true birth rates $(b_1, b_2) = (0.8, 0.5)$, known death rates $(d_1, d_2) = (0.1, 0.1)$, and known switching rates $(s_{12}, s_{21}) = (0.6, 0.4)$, all per cell per day. Only the birth rates are fitted, over the region $(b_1, b_2) \in [0.3, 1.1]^2$. The experiments without death set $d_1 = d_2 = 0$ and use the same true birth rates, switching rates, and fitting region. At the true rates in the birth–death experiment, $L = 0.542443$ and $\lambda_2 = -0.442443$ per day. Throughout the birth–death fitting region, $L \geq 0.2$ per day. Hypothesis (H4), $\lambda_2 < L/2$ for the two-type model, follows from

$$\frac{L}{2} - \lambda_2 = \frac{3\sqrt{(A_{11} - A_{22})^2 + 4s_{12}s_{21}} - (A_{11} + A_{22})}{4}. \tag{S19}$$

With $s_{12} = 0.6$, $s_{21} = 0.4$, and $A_{11} + A_{22} \le 1$, this expression is strictly positive throughout the birth–death fitting region. It remains positive in the experiments without death, where $A_{11} + A_{22} \le 1.2$. Irreducibility and the eigenvalue condition, hypotheses (H2) and (H4) of the main text, therefore hold over both fitting regions, not just at the true rates.

#### S2.2 Extinction and survival probabilities

Let $\zeta_i(t)$ be the probability that the population is extinct by time $t$, starting from one type-$i$ cell, and let $\zeta_i(\infty) = \lim_{t\to\infty} \zeta_i(t)$ be its eventual extinction probability. Conditioning on the first event gives the backward equations (Athreya and Ney 1972, Chapter V, Section 7.1, Equation (2)), for $i, j \in \{1, 2\}$ with $j \ne i$,

$$\zeta_i'(t) = d_i - (b_i + d_i + s_{ij})\zeta_i(t) + b_i\zeta_i(t)^2 + s_{ij}\zeta_j(t), \qquad \zeta_i(0) = 0, \tag{S20}$$

where the prime denotes differentiation with respect to time $t$. The term $b_i\zeta_i(t)^2$ arises because, after division, the two daughter lineages evolve independently and both must become extinct. The eventual probabilities $\zeta_1(\infty)$ and $\zeta_2(\infty)$ are the smallest nonnegative stationary solution, obtained by monotone iteration from zero. Independent lineages give the probabilities $\pi_t$ of survival to time $t$ and $\pi_\infty$ of eventual survival,

$$\pi_t = 1 - \zeta_1(t)^{n_1}\zeta_2(t)^{n_2}, \qquad \pi_\infty = 1 - \zeta_1(\infty)^{n_1}\zeta_2(\infty)^{n_2}. \tag{S21}$$

At the true rates, the one-founder eventual extinction probabilities $\zeta_1(\infty)$ and $\zeta_2(\infty)$ are about 0.148 and 0.174. The logarithms of the extinction and survival probabilities are stored separately, so that an extremely small but positive extinction weight is retained even when the survival probability rounds to one.

#### S2.3 Error from the eventual-survival substitution

Among populations alive at time $t$, some survive forever and others eventually become extinct. Write $\mathcal{A}_t$ for survival at time $t$, $\mathcal{S}$ for eventual survival, and $\mathcal{S}^c$ for eventual extinction. Since $\mathcal{S} \subseteq \mathcal{A}_t$, their conditional probabilities are

$$\Pr(\mathcal{S} \mid \mathcal{A}_t) = \frac{\pi_\infty}{\pi_t}, \qquad \varepsilon_t = \Pr(\mathcal{S}^c \mid \mathcal{A}_t) = 1 - \frac{\pi_\infty}{\pi_t},$$

as in Equation (17) of the main text. The observation of a culture at time $t$ is $Y = Z(t) + \epsilon$, where the measurement error $\epsilon \sim \mathcal{N}_2(0, R)$ is independent of the population (Equation (9) of the main text). In the experiments, $R = 4I_2$. For an event $\mathcal{E}$, write $\Pr(Y \in \cdot \mid \mathcal{E})$ for the law of $Y$ conditional on $\mathcal{E}$. By the law of total probability, applied to each measurable set, the observation law conditional on survival at time $t$ satisfies

$$\Pr(Y \in \cdot \mid \mathcal{A}_t) = (1 - \varepsilon_t)\Pr(Y \in \cdot \mid \mathcal{S}) + \varepsilon_t \Pr(Y \in \cdot \mid \mathcal{A}_t \cap \mathcal{S}^c). \tag{S22}$$

If $\varepsilon_t = 0$, the last term is omitted and the two survival-conditioned laws coincide. The fraction of populations alive at time $t$ that eventually become extinct is $\varepsilon_t$. By Equation (S22), replacing conditioning on survival at time $t$ by conditioning on eventual survival changes the observation law by at most $\varepsilon_t$ in total variation distance:

$$d_{\mathrm{TV}}(\Pr(Y \in \cdot \mid \mathcal{A}_t), \Pr(Y \in \cdot \mid \mathcal{S})) \le \varepsilon_t, \tag{S23}$$

where $d_{\mathrm{TV}}$ denotes total variation distance: the supremum, over measurable events, of the absolute difference between the probabilities assigned by the two laws. The argument applies to the observation $Y$, including the independent measurement noise. In the observation law for a seeded culture, the law conditional on survival at time $t$ has weight $\pi_t$. Keeping this weight and the law conditional on extinction at time $t$ unchanged, the eventual-survival substitution alone contributes at most $\pi_t \varepsilon_t = \pi_t - \pi_\infty$ in total variation distance.

Averaging $\widehat{\mu}(t, w) = wm(t)/a_0$ over the distribution of the positive amplitude gives the approximate mean $m(t)/\pi_\infty$ for the living-culture component, because the mean of the positive amplitude is $a_0/\pi_\infty$. The exact mean conditional on survival at time $t$ is $m/\pi_t$. Consequently, the mean of the full observation density of $M$ is $\pi_t m/\pi_\infty$. No additional mean correction is applied.

For $G$, $EG$, $M$, and $M_0$ in the birth–death experiment, the largest recorded $\varepsilon_t$ over all fitted rate vectors is 0.0037443. Over the rate vectors at the boundaries of the profile-likelihood confidence sets, the largest value is 0.0050885. For each founder count, we check both founder types and both observation times. At the lower corner $(0.3, 0.3)$, the one-founder values are 0.05510 and 0.03693 at days 9 and 11. Small values of $\varepsilon_t$ at fitted rates do not establish a uniformly small error bound for the eventual-survival substitution over the entire fitting region.

### S3 Exact Yule reference distribution

The Yule process $N(t)$ starts from one cell, and each cell divides at rate $b$ without death or switching. Figure 1 of the main text compares its approximate and exact distributions at days 0.5 and 5, with $b = 0.8$ per day. Its count at time $t$ is geometric on the positive integers (Kendall 1948), with probabilities

$$p_j = \Pr(N(t) = j) = \exp(-bt)(1 - \exp(-bt))^{j-1}, \qquad j \geq 1.$$

For an observation $Y = N(t) + \epsilon$ with independent measurement error $\epsilon \sim \mathcal{N}(0, 4)$, the exact density and cumulative distribution function at an observed value $x$ are

$$p_{\text{exact}}(x) = \sum_{j \geq 1} p_j \varphi_1(x; j, 4), \qquad F_{\text{exact}}(x) = \sum_{j \geq 1} p_j \Phi((x - j)/2), \tag{S24}$$

where $\varphi_1$ is a univariate Gaussian density and $\Phi$ the standard normal cumulative distribution function. For numerical evaluation, the infinite sums are truncated so that the omitted count probability is at most $10^{-12}$ in the reported calculations and $10^{-14}$ in the numerical accuracy checks. The omitted probability is the geometric tail $\Pr(N(t) > J) = (1 - \exp(-bt))^J$ beyond the last retained count $J$, and the retained probabilities are not renormalized.

The approximations follow Section 2.3 of the main text with $a_0 = 1$. The founder-aware likelihood $M$ integrates Gaussian distributions with mean $\widehat{\mu}(t, w) = w \exp(bt)$ and variance $\widehat{Q}(t, w) = w \exp(bt)$ over the distribution of $W$. Here, $W$ has an exact exponential distribution with mean one. The variant $M_0$ sets $\widehat{Q} = 0$. The population Gaussian $G$ uses the exact mean $\exp(bt)$ and variance $\exp(2bt) - \exp(bt)$ of Equation (5) of the main text. Without death, no culture becomes empty, so $\pi_t = 1$ and $G$ and $EG$ coincide.

Quantiles use the generalized inverse $F^{-1}(u) = \inf\{x : F(x) \geq u\}$. Figure 1 of the main text uses $u = 0.001, 0.01, 0.025, 0.05, 0.1, 0.25, 0.5, 0.75, 0.9, 0.95, 0.975, 0.99, 0.999$ for every likelihood. For comparisons of cell counts, the reference quantiles remain integer-valued, and negative quantiles from the Gaussian approximations are retained.

### S4 Numerical methods and accuracy checks

#### S4.1 Likelihood evaluation and parameter estimation

Each dataset contains 40 independent cultures with the same founder count, either one or 50. There are ten cultures for each combination of founder type and observation time, with all founders of a culture belonging to the same type. The birth–death experiment uses days 9 and 11; the two experiments without death use days 5 and 7 and days 9 and 11, respectively. Each experiment has 100 datasets per founder count, and the likelihoods being compared are fitted to the same observations. For each culture, the observation is the vector of counts plus independent Gaussian measurement error with covariance $R = 4I_2$, as in Section S2.3. All cultures are retained, including empty cultures, and inference uses the noisy observations without access to simulated counts or survival status.

To evaluate the living-culture component of each mixture likelihood, we integrate over the positive establishment amplitude using $u = \log w$, including the Jacobian $dw = \exp(u)\,du$. The integration intervals are chosen to include peaks of the integrand. On each interval, we compare 16-point and 32-point Gauss–Legendre estimates of the integral. We bisect the interval when their difference exceeds the prescribed tolerance. No lower bound is imposed on computed densities, and the truncated amplitude distribution is not renormalized. Numerical settings are listed with the accuracy checks in Table S1. The accompanying code and experiment configurations specify the search grids, starting points, and random seeds.

Optimization combines a grid search, refinement from several starting points, explicit boundary checks, and bounded Nelder–Mead optimization. For each profile, we fix one birth rate and reoptimize the other fitted parameters at every value examined, including each boundary of the profile-likelihood confidence set, or profile set. As defined in Equation (20) of the main text, the nominal 90% profile set contains the values for which twice the decrease from the maximized log likelihood is at most 2.705543, the 90th percentile of a chi-square distribution with one degree of freedom. The search allows disconnected profile sets and sets truncated by the fitting region, although a finite grid cannot rule out arbitrarily narrow missed components.

For $M_{\text{const}}$, the search includes $v = 0$ and uses the transformed variance $\log(1+v/4)$, where 4 is the known measurement variance. The model allows arbitrarily large $v$, but the numerical search requires a finite interval $0 \le v \le \Omega$. We choose this limit adaptively, using a bound on the log likelihood to check whether larger variances could improve the fit. Under $M_{\text{const}}$, each Gaussian in the living-culture component has covariance $vI_2 + R = (4+v)I_2$, so for $v \ge \Omega$ its density is at most $[2\pi(4+\Omega)]^{-1}$. Let $\varphi_2(y;\mu,C)$ denote the bivariate Gaussian density at $y$ with mean $\mu$ and covariance $C$, and let $k_0(y) = \varphi_2(y;0,R)$ be the observation density of an empty culture, as in Equation (16) of the main text. The observation density $p_{M_{\text{const}}}$ is a weighted average of the living-culture density and $k_0$, so it is at most the larger of the two. Let $y_1,\dots,y_{40}$ be the observations of one dataset, and let $\theta = (b_1, b_2)$ be the fitted birth rates, as in the main text. The log likelihood of the dataset is $\mathcal{L}_{M_{\text{const}}}(\theta,v) = \sum_{j=1}^{40} \log p_{M_{\text{const}}}(y_j;\theta,v)$. For every $\theta$ in the fitting region and every $v \ge \Omega$, it therefore satisfies

$$\mathcal{L}_{M_{\text{const}}}(\theta,v) \le \mathcal{U}(\Omega) = \sum_{j=1}^{40} \max\{\log k_0(y_j), -\log[2\pi(4+\Omega)]\}. \tag{S25}$$

The bound $\mathcal{U}(\Omega)$ depends on neither $\theta$ nor $v$, so it bounds the log likelihood over all variances $v \ge \Omega$. The initial limit is the larger of the measurement variance and the mean of the squared components of the observations. After searching within the current limit, we compare $\mathcal{U}(\Omega)$ with the best log likelihood found, evaluated with both the usual and tighter integration settings. If $\mathcal{U}(\Omega)$ is not below the log likelihood computed with each integration setting by the required numerical margin, we increase $\Omega$ tenfold and repeat the search. For each profile calculation, the birth rate being profiled is held fixed throughout the search and comparison. This procedure restricts the search to a finite interval while checking that no variance beyond the accepted limit can give a larger log likelihood, subject to numerical accuracy.

#### S4.2 Verification of the calculations

The checks below assess the simulator and the numerical calculation of the specified likelihoods, estimates, and profile sets. They do not establish the accuracy of the approximate likelihoods as descriptions of the branching observation law.

The exact event simulation of Gillespie (1977) was checked on cultures generated independently of the inference and predictive samples. For each founder type and observation time, the check used 10,000 cultures with one founder and 500 with 50 founders. Sample means and covariance entries differed from their exact values by at most 2.3 estimated Monte Carlo standard errors, and extinction frequencies passed the prespecified binomial checks. No simulated population reached the computational size limit, and no culture was discarded or replaced.

To evaluate the likelihoods, we compute the population mean $m(t)$ and covariance $V(t)$ from the moment equations $m' = Am$, with $m(0) = z_0$, and $V' = AV + VA^\top + D(m)$, with $V(0) = 0$ (Section 2.2 of the main text). Because $D$ is linear, $m$ and the entries of $V$ together satisfy one linear system of differential equations with constant coefficients, which we solve with a single matrix exponential rather than by numerical time stepping. The covariance $V(t)$ used by $G$ and $EG$ was checked by independently evaluating the formula of Gunnarsson et al. (2023, Section 3.3 and Appendix A.2), after transposing their row-vector convention to our column-vector convention. At the true rates and both observation times, the relative difference from the calculation using the moment equations was below $10^{-12}$ in Frobenius norm.

Adding the approximate conditional covariance $\widehat{Q}(t, w)$ to the measurement covariance $R$ cannot reduce the variance in any direction. Consequently, omitting probability mass $\varepsilon_{\text{tail}}$ from the distribution of the positive amplitude contributes at most $\varepsilon_{\text{tail}}/(2\pi\sqrt{\det R})$ to the error in a two-type density. This absolute bound need not be small relative to the density, so we also check changes in log density. Observation densities were reevaluated using the log-amplitude integration of Section S4.1 with tighter tolerances and smaller omitted amplitude tails, as specified in Table S1. An additional check integrates directly in $w$ using adaptive 21-point Gauss–Kronrod integration with extrapolation (QUADPACK (Piessens et al. 1983), through SciPy's `quad` function (Virtanen et al. 2020)). This calculation uses the same conditional density and tail limits. These checks include near-zero and negative observations, corners of the fitting region, and representative fitted parameters. We check the density of the living-culture component separately from the full observation density, so that the empty-culture component does not mask errors in integration over amplitude. The first row of Table S2 reports the largest changes in log density across these checks.

Table S1: Numerical settings for the birth–death experiment and its accuracy checks. Tolerance pairs are relative and absolute tolerances, respectively. The interval for integration over amplitude is expanded as needed to include peaks associated with each observation.

| Setting | Reported results | Accuracy checks |
|---|---|---|
| Tolerances for integration over amplitude | $2 \times 10^{-7}$, $10^{-10}$ | $2 \times 10^{-10}$, $10^{-13}$ |
| Initial omitted probability in each amplitude tail | $10^{-10}$ | $10^{-12}$ |
| Tolerances for extinction equations | $10^{-10}$, $10^{-12}$ | $10^{-12}$, $10^{-14}$ |
| Tolerance for profile-set boundaries (per day) | $10^{-5}$ | $10^{-6}$ |

We repeated fits and profiles for all five likelihoods on the first five datasets at each founder count, which were selected independently of the results. These calculations used tighter numerical tolerances, denser grids, and more starting points. The remaining rows of Table S2 report the largest changes in fitted rates, maximized log likelihoods, and profile-set boundaries.

Under the tighter settings, the profile sets had the same number of components, the same truncation status, and the same coverage. A further check of the set whose boundary was

Table S2: Largest numerical changes under tighter settings or alternative integration in the birth–death experiment. Changes in log density are dimensionless; changes in estimates and set boundaries are per day.

| Quantity checked | $G$, $EG$, $M$, $M_0$ | $M_{\text{const}}$ |
|---|---|---|
| Observation log density, tighter or independent integration | $3.79 \times 10^{-9}$ | $9.78 \times 10^{-10}$ |
| Fitted birth rate, tighter refit | $4.43 \times 10^{-8}$ | $5.64 \times 10^{-9}$ |
| Maximized log likelihood of a dataset, tighter refit | $4.51 \times 10^{-10}$ | $3.39 \times 10^{-10}$ |
| Profile-set boundary, tighter search | $2.53 \times 10^{-6}$ | $2.44 \times 10^{-6}$ |

Table S3: Computational times for the birth–death experiment. Fit times are means over five datasets per founder count $N_0$; profile times cover both birth rates of one representative dataset per founder count. Evaluation times are for the log likelihood of one dataset with 40 observations. A dash denotes a timing not tabulated here. For $M_{\text{const}}$, the first value excludes time spent saving intermediate search results, as in the main-text ratios; parentheses give the total elapsed time. For $M$, only completed fits and profiles are saved.

| Likelihood | $N_0$ | Fit (s) | Evaluation (ms) | Profiles (s) |
|---|---|---|---|---|
| Comparison with Gaussian likelihoods | | | | |
| $G$ | 1 | 0.857 | 1.29 | 12.5 |
| $G$ | 50 | 0.984 | 1.21 | 10.2 |
| $EG$ | 1 | 5.09 | — | — |
| $EG$ | 50 | 5.02 | — | — |
| $M$ | 1 | 5.60 | 4.23 | 68.1 |
| $M$ | 50 | 4.90 | 4.26 | 67.6 |
| Comparison with a fitted constant conditional variance | | | | |
| $M$ | 1 | 1.44 | — | 20.1 |
| $M$ | 50 | 1.43 | — | 19.1 |
| $M_{\text{const}}$ | 1 | 7.45 (9.43) | — | 451 (562) |
| $M_{\text{const}}$ | 50 | 10.10 (12.52) | — | 402 (512) |

closest to a true rate also left its coverage unchanged. All fits and profiles completed, and all profile sets were connected and contained within the fitting region. Some local optimization attempts failed: nine during fitting of $G$, $EG$, $M$, and $M_0$ combined, and ten during fitting and 324 during profiling of $M_{\text{const}}$. Other starting points completed each search successfully.

For the Yule reference calculations, tighter truncation and integration changed quantiles by less than $10^{-5}$ cells, much less than the quantile discrepancies in Figure 1 of the main text.

Both experiments without death also passed checks of moments, densities, fits, and profiles.

##### S4.3 Computational cost

Table S3 gives the absolute computation times used to calculate the ratios reported in the main text. Times include all calculations required by each likelihood; extinction probabilities, for example, are not computed for $G$. The comparison with the Gaussian likelihoods used four concurrent processes with one numerical thread each. The separate comparison with $M_{\text{const}}$ used one process with one thread and a shared compiled solver for extinction probabilities. Times should therefore be compared within each comparison only. Previously computed quantities were cleared before each fit and profile calculation; individual likelihood evaluations were timed after reusable quantities had been computed.

In the comparison with the Gaussian likelihoods, extinction calculations accounted for about three fifths of the fitting time under $M$, and amplitude integration for about one fifth.

#### S5 Additional simulation results

Table S4 gives the differences in log density shown in Figure 2 of the main text, including the comparison with $M_0$ that is omitted from the figure. Tables S5 and S6 complement the birth–death inference results in Figure 3 of the main text. They report paired differences, absolute root-mean-square errors (RMSEs), and signed biases, using all datasets. For paired comparisons, we estimate uncertainty by resampling whole datasets, so that each bootstrap resample compares the likelihoods on the same observations. For each RMSE reduction, both RMSEs are recomputed across the resampled datasets before forming their ratio. The paired bootstrap uses 2,000 resamples for the birth–death experiment and 5,000 for the two experiments without death. Table S7 reports the one-founder inference results for the two experiments without death.

Table S4: Mean paired log-density differences $\log p_M(Y) - \log p_H(Y)$ for comparator $H$ at the true birth–death rates, with Monte Carlo standard errors in parentheses. Each entry uses 500 independent observations for the indicated founder count $N_0$, pure founder type, and observation day. All likelihoods are evaluated on the same cultures. Positive values favor $M$.

| $N_0$ | Founder type | Day | $M$ minus $G$ | $M$ minus $EG$ | $M$ minus $M_0$ |
|---|---|---|---|---|---|
| 1 | 1 | 9 | 1.163 (0.075) | 0.553 (0.043) | 13.6 (1.7) |
| 1 | 1 | 11 | 1.152 (0.087) | 0.570 (0.047) | 48.3 (4.9) |
| 1 | 2 | 9 | 1.024 (0.079) | 0.487 (0.049) | 12.0 (1.5) |
| 1 | 2 | 11 | 1.506 (0.126) | 0.734 (0.087) | 50.9 (7.7) |
| 50 | 1 | 9 | 0.015 (0.008) | 0.015 (0.008) | 933.9 (58.2) |
| 50 | 1 | 11 | 0.024 (0.009) | 0.024 (0.009) | 2460.9 (155.9) |
| 50 | 2 | 9 | 0.019 (0.009) | 0.019 (0.009) | 625.6 (39.0) |
| 50 | 2 | 11 | 0.008 (0.008) | 0.008 (0.008) | 1938.5 (135.0) |

Table S5: Paired comparisons of $M$ with each comparator in the birth–death experiment, with 95% bootstrap intervals. RMSE reduction is $100(1 - \text{RMSE}_M/\text{RMSE}_H)$ for comparator $H$; positive values favor $M$. Coverage gains are in percentage points (pp), and changes in mean total set length are in $10^{-3}$ per day. Length changes are absolute differences, whereas the main text reports percentage increases. All sets use the common unadjusted chi-square rule, so coverage gains must be assessed with their associated length changes.

| $N_0$ | Rate | Comparator | RMSE reduction (%) | Coverage gain (pp) | Length change |
|---|---|---|---|---|---|
| 1 | $b_1$ | $G$ | 16.7 [9.9, 22.9] | 34.0 [25.0, 43.0] | 35.15 [34.69, 35.63] |
| 1 | $b_1$ | $EG$ | 16.4 [9.4, 23.1] | 33.0 [24.0, 42.0] | 31.82 [31.50, 32.18] |
| 1 | $b_1$ | $M_0$ | 6.6 [−0.8, 13.6] | 8.0 [2.0, 14.0] | 11.43 [10.97, 11.90] |
| 1 | $b_1$ | $M_{\text{const}}$ | 6.2 [−1.5, 13.4] | 6.0 [1.0, 12.0] | 5.99 [5.53, 6.47] |
| 1 | $b_2$ | $G$ | 8.1 [−3.5, 18.4] | 30.0 [21.0, 40.0] | 36.72 [36.06, 37.38] |
| 1 | $b_2$ | $EG$ | 7.3 [−5.9, 17.3] | 25.0 [17.0, 34.0] | 32.71 [32.22, 33.23] |
| 1 | $b_2$ | $M_0$ | −2.5 [−12.0, 5.8] | 9.0 [3.0, 15.0] | 16.84 [16.19, 17.48] |
| 1 | $b_2$ | $M_{\text{const}}$ | −5.3 [−15.5, 3.8] | 4.0 [−1.0, 10.0] | 9.55 [8.87, 10.20] |
| 50 | $b_1$ | $G$ | 0.0 [−4.2, 4.0] | 2.0 [−2.0, 6.0] | 0.18 [0.16, 0.19] |
| 50 | $b_1$ | $EG$ | 0.0 [−4.3, 4.3] | 2.0 [−2.0, 6.0] | 0.18 [0.16, 0.19] |
| 50 | $b_1$ | $M_0$ | −0.1 [−3.7, 3.7] | 8.0 [3.0, 14.0] | 1.59 [1.58, 1.60] |
| 50 | $b_1$ | $M_{\text{const}}$ | −0.4 [−3.9, 3.2] | 4.0 [1.0, 8.0] | 0.34 [0.28, 0.40] |
| 50 | $b_2$ | $G$ | −1.1 [−6.1, 3.7] | 0.0 [−4.0, 4.0] | 0.59 [0.58, 0.60] |
| 50 | $b_2$ | $EG$ | −1.1 [−6.2, 3.6] | 0.0 [−4.0, 4.0] | 0.59 [0.58, 0.60] |
| 50 | $b_2$ | $M_0$ | 2.5 [−2.4, 7.1] | 11.0 [5.0, 18.0] | 2.25 [2.24, 2.26] |
| 50 | $b_2$ | $M_{\text{const}}$ | 2.6 [−2.4, 7.0] | 0.0 [−4.0, 4.0] | 0.60 [0.52, 0.68] |

Table S6: Signed birth-rate bias and absolute RMSE, in inverse days, over 100 datasets per founder count. Parentheses give the Monte Carlo standard error (MCSE) of bias; brackets give individual 95% bootstrap intervals for RMSE. The likelihoods share observations within every dataset. The paired RMSE comparisons appear in Table S5 and the main-text inference figure.

| $N_0$ | Rate | Method | Bias (MCSE) | RMSE [95% interval] |
|---|---|---|---|---|
| 1 | $b_1$ | $G$ | −0.00448 (0.00294) | 0.02960 [0.02537, 0.03425] |
| 1 | $b_1$ | $EG$ | −0.00365 (0.00294) | 0.02948 [0.02519, 0.03387] |
| 1 | $b_1$ | $M_0$ | −0.00163 (0.00265) | 0.02640 [0.02217, 0.03041] |
| 1 | $b_1$ | $M_{\mathrm{const}}$ | −0.00018 (0.00264) | 0.02628 [0.02214, 0.03026] |
| 1 | $b_1$ | $M$ | −0.00272 (0.00246) | 0.02465 [0.02036, 0.02894] |
| 1 | $b_2$ | $G$ | −0.00616 (0.00258) | 0.02638 [0.02270, 0.03035] |
| 1 | $b_2$ | $EG$ | −0.00522 (0.00258) | 0.02615 [0.02234, 0.03001] |
| 1 | $b_2$ | $M_0$ | −0.00448 (0.00233) | 0.02366 [0.02028, 0.02690] |
| 1 | $b_2$ | $M_{\mathrm{const}}$ | −0.00239 (0.00230) | 0.02302 [0.01993, 0.02602] |
| 1 | $b_2$ | $M$ | −0.00347 (0.00241) | 0.02424 [0.02103, 0.02725] |
| 50 | $b_1$ | $G$ | 0.00073 (0.00037) | 0.00372 [0.00320, 0.00422] |
| 50 | $b_1$ | $EG$ | 0.00073 (0.00037) | 0.00372 [0.00320, 0.00422] |
| 50 | $b_1$ | $M_0$ | 0.00100 (0.00036) | 0.00372 [0.00318, 0.00423] |
| 50 | $b_1$ | $M_{\mathrm{const}}$ | 0.00098 (0.00036) | 0.00371 [0.00320, 0.00421] |
| 50 | $b_1$ | $M$ | 0.00104 (0.00036) | 0.00372 [0.00320, 0.00426] |
| 50 | $b_2$ | $G$ | 0.00029 (0.00030) | 0.00301 [0.00264, 0.00338] |
| 50 | $b_2$ | $EG$ | 0.00029 (0.00030) | 0.00301 [0.00263, 0.00338] |
| 50 | $b_2$ | $M_0$ | 0.00055 (0.00031) | 0.00312 [0.00273, 0.00351] |
| 50 | $b_2$ | $M_{\mathrm{const}}$ | 0.00057 (0.00031) | 0.00312 [0.00274, 0.00350] |
| 50 | $b_2$ | $M$ | 0.00049 (0.00030) | 0.00304 [0.00269, 0.00339] |

Table S7: Experiments without death, with one founder per culture and 100 datasets for each window of observation days. The two comparison columns give percentage RMSE reductions for $M$ with paired 95% bootstrap intervals. Coverage percentages are listed in the order $G/M/M_0$ for nominal 90% sets. Length ratio is mean length under $M$ divided by that under $G$. The windows use independent datasets.

| Days | Rate | vs. $G$ | vs. $M_0$ | Coverage | Length ratio |
|---|---|---|---|---|---|
| 5, 7 | $b_1$ | 6.6 [−1.3, 13.3] | 0.8 [−7.3, 8.8] | 64/89/67 | 1.71 |
| 5, 7 | $b_2$ | 18.5 [10.5, 25.5] | 19.3 [11.0, 26.8] | 64/95/73 | 1.63 |
| 9, 11 | $b_1$ | 9.1 [0.8, 16.9] | 2.8 [−3.1, 9.2] | 58/87/84 | 1.87 |
| 9, 11 | $b_2$ | 14.3 [6.6, 20.9] | 12.0 [7.9, 16.6] | 52/86/75 | 1.82 |

### References


Asmussen, S., Hering, H.: Branching Processes. Progress in Probability and Statistics, vol. 3. Birkhäuser, Boston (1983). https://doi.org/10.1007/978-1-4615-8155-0

Athreya, K.B., Ney, P.E.: Branching Processes. Springer, Berlin (1972). https://doi.org/10.1007/978-3-642-65371-1

Gunnarsson, E.B., Foo, J., Leder, K.: Statistical inference of the rates of cell proliferation and phenotypic switching in cancer. Journal of Theoretical Biology **568**, 111497 (2023) https://doi.org/10.1016/j.jtbi.2023.111497

Gillespie, D.T.: Exact stochastic simulation of coupled chemical reactions. The Journal of Physical Chemistry **81**(25), 2340–2361 (1977) https://doi.org/10.1021/j100540a008

Janson, S.: Functional limit theorems for multitype branching processes and generalized Pólya urns. Stochastic Processes and their Applications **110**, 177–245 (2004) https://doi.org/10.1016/j.spa.2003.12.002

Kendall, D.G.: On the generalized "birth-and-death" process. The Annals of Mathematical Statistics **19**(1), 1–15 (1948) https://doi.org/10.1214/aoms/1177730285

Kolesko, K., Kołodziejska, A., Meiners, M.: Asymptotic fluctuations in supercritical multi-type Crump–Mode–Jagers processes. arXiv preprint arXiv:2607.22414v1 (2026). https://arxiv.org/abs/2607.22414v1

Piessens, R., Doncker-Kapenga, E., Überhuber, C.W., Kahaner, D.K.: QUADPACK: A Subroutine Package for Automatic Integration. Springer Series in Computational Mathematics, vol. 1. Springer, Berlin, Heidelberg (1983). https://doi.org/10.1007/978-3-642-61786-7

Protter, P.E.: Stochastic Integration and Differential Equations, 2nd edn. Springer, Berlin, Heidelberg (2005). https://doi.org/10.1007/978-3-662-10061-5 . Version 2.1, corrected third printing

Ren, Y.-X., Song, R., Zhang, R.: Central limit theorems for supercritical branching Markov processes. Journal of Functional Analysis **266**(3), 1716–1756 (2014) https://doi.org/10.1016/j.jfa.2013.10.015

Ren, Y.-X., Song, R., Zhang, R.: Central limit theorems for supercritical branching nonsymmetric Markov processes. Annals of Probability **45**(1), 564–623 (2017) https://doi.org/10.1214/14-AOP987

Virtanen, P., Gommers, R., Oliphant, T.E., Haberland, M., Reddy, T., Cournapeau, D., Burovski, E., Peterson, P., Weckesser, W., Bright, J., Walt, S.J., Brett, M., *et al.*: SciPy 1.0: fundamental algorithms for scientific computing in Python. Nature Methods **17**(3), 261–272 (2020) https://doi.org/10.1038/s41592-019-0686-2
